# CF-Seq, An Accessible Web Application for Rapid Re-Analysis of Cystic Fibrosis Pathogen RNA Sequencing Studies

**DOI:** 10.1101/2022.03.07.483313

**Authors:** Samuel L. Neff, Thomas H. Hampton, Charles Puerner, Liviu Cengher, Georgia Doing, Alexandra J. Lee, Katja Koeppen, Ambrose L. Cheung, Deborah A. Hogan, Robert A. Cramer, Bruce A. Stanton

## Abstract

Researchers studying cystic fibrosis (CF) pathogens have produced numerous RNA-seq datasets which are available in the gene expression omnibus (GEO). Although these studies are publicly available, substantial computational expertise and manual effort are required to compare similar studies, visualize gene expression patterns within studies, and use published data to generate new experimental hypotheses. Furthermore, it is difficult to filter available studies by domain-relevant attributes such as strain, treatment, or media, or for a researcher to assess how a specific gene responds to various experimental conditions across studies. To reduce these barriers to data re-analysis, we have developed an R Shiny application called CF-Seq, which works with a compendium of 147 studies and 1,446 individual samples from 13 clinically relevant CF pathogens. The application allows users to filter studies by experimental factors and to view complex differential gene expression analyses at the click of a button. Here we present a series of use cases that demonstrate the application is a useful and efficient tool for new hypothesis generation. (CFSeq: http://scangeo.dartmouth.edu/CFSeq/)

## Introduction

Cystic fibrosis (CF) is a monogenic, homozygous recessive genetic disease that affects over 30,000 people in the US and more than 70,000 worldwide^1^. The disease is caused by mutations of the cystic fibrosis transmembrane conductance regulator (CFTR) gene, which is expressed in a wide variety of cells throughout the body but has been predominantly studied in the context of the lungs and the digestive system^2,3,4,5^. In the lungs, the absence of CFTR contributes to mucus obstruction, chronic microbial infections, systemic inflammation, and progressive lung disease, which is the leading cause of mortality^6,7,8,9^. Furthermore, people with CF (pwCF) are commonly diagnosed with exocrine pancreatic insufficiency, and tend to exhibit microbial dysbiosis in the GI tract, which both contribute to nutritional deficits, poor growth, and a myriad of other GI symptoms^5,10,11^. Based on population data from the Cystic Fibrosis Foundation Patient Registry, pwCF born between 2015 and 2019 have a median life expectancy of just 46 years^12^. The life expectancy for pwCF has risen dramatically over the past few decades, however, as scientists, pharmaceutical companies, and physicians have developed new drugs to treat the molecular defect in CF, standardized clinical guidelines, and produced new antibiotic regimens to manage persistent bacterial infections^13^.

Given the contribution of invasive pathogens to lung disease progression, lung microbiology has long been a key focus of CF research. CF researchers have traditionally studied a suite of “classic CF pathogens’’ that are known to infect the CF lungs and exacerbate lung disease. These pathogens include the gram-negative bacterium *Pseudomonas aeruginosa*, the gram-positive bacterium *Staphylococcus aureus*, gram-negative bacteria of the genus *Burkholderia*, and fungal species such as *Aspergillus fumigatus* [Table 1]. In recent years, the set of recognized CF pathogens has expanded as epidemiological studies have identified species that are rising in prevalence and impacting clinical outcomes (e.g., non-tuberculous mycobacteria species such as *M. abscessus*)^14,15^. In addition, more sensitive culture tools have allowed researchers to recognize the clinical relevance of less prevalent aerobic and anaerobic species^16,17^. Recently, researchers have begun to develop model systems to interrogate the interactions between CF pathogens in the lungs and to consider how the overall shape of the CF community – the diversity and abundance of different bacteria – contributes to clinical outcomes^18^. In fact, studies have found that a patient’s microbial community as a whole may be more effective at predicting disease outcomes than colonization with any individual species^19^.

**Table 1.** Clinical relevance of cystic fibrosis pathogens with studies featured in the CF-Seq application

| Genus | Species | Type | Location | Clinical Relevance |
| --- | --- | --- | --- | --- |
| *Aspergillus* | *Aspergillus fumigatus* | Fungus | Lung | - *Aspergillus* colonization is associated with lower lung function and more frequent hospitalization<sup>64</sup><br>- Long-term colonization is also associated with declining chest computed tomography (CT) scores over time<sup>65</sup><br>- *Aspergillus* colonization can result in a hard-to-treat chronic condition called allergic bronchopulmonary aspergillosis (ABPA)<sup>66</sup> |
| *Bacteroides* | Multiple | Bacterium | Gut | - For infants with CF, the stool microbiota does not exhibit the increase in diversity that is observed in infants without CF over the first several years of life. Specifically, infants with CF tend to see reduced levels of the genus *Bacteroides* (the same has been noted for *Veillonella* species, *Prevotella* species, and *Bifidobacterium* species as well)<sup>67,68</sup><br>- Exposure of intestinal cell lines to *Bacteroides* species supernatant has the effect of reducing IL-8 production, suggesting that absence of *Bacteroides* species may contribute to inflammation in the CF gut<sup>67</sup> |
| *Burkholderia* | Multiple | Bacterium | Lung | - *Burkholderia* species infections are extremely difficult to eradicate after<br>the bacteria occupy the CF lungs and establishes chronic infection<sup>69</sup><br>- Associated with highly unpredictable clinical symptoms, in some cases severe decline in lung function and high mortality<sup>69</sup><br>- *Burkholderia cenocepacia* and *Burkholderia multivorans* are the most common species isolated from the CF lungs, with *B. cenocepacia* infections generally being more severe<sup>69</sup><br>- Like *Pseudomonas*, *Burkholderia* can be found as a dominant pathogen in the lungs of some adult pwCF<sup>70</sup><br>- *Burkholderia* infection is a common contraindication for lung transplant<sup>71</sup> |
| *Candida* | *Candida albicans* | Fungus | Lung | - *Candida albicans* is the most common fungus isolated from the lungs of pwCF<sup>72</sup><br>- pwCF receiving frequent IV antibiotics are susceptible to *Candida* sepsis, requiring extensive anti-fungal treatment and precluding further IV antibiotic treatment<sup>72</sup><br>- A mild association between *Candida albicans* colonization and lung function decline has been observed<sup>73</sup> |
| *Clostridium* | *Clostridium difficile* | Bacterium | Gut | - pwCF present with a much higher prevalence of GI colonization with *C. difficile* than the general population, although<br>severe *C. difficile* infection is not common and carriage of *C. difficile* is usually asymptomatic<sup>74</sup><br>- High prevalence of *C. difficile* in the CF population is thought to stem from frequent antibiotic use and hospital-based exposure<sup>75</sup> |
| *Fusobacterium* | *Fusobacterium nucleatum* | Bacterium | Lung | - *Fusobacterium nucleatum*, an anaerobe, is known to be present on occasion in the cystic fibrosis lung, with one study finding that of 109 pwCF, it was detected in 5.5% of subjects<sup>76</sup><br>- A study has shown that *F. nucleatum* and *P. aeruginosa* frequently coexist in the lungs – and this co-culture state has been found to be associated with increased bacterial count and bacterial tolerance to antibiotics<sup>77</sup><br>- Production of short-chain fatty acids by anaerobes has been found to induce IL-8 production in CF bronchial epithelial cells in vitro, which may contribute to pulmonary inflammation in vivo<sup>76</sup> |
| *Haemophilus* | *Haemophilus influenza* | Bacterium | Lung | - *H. influenza* commonly infects pwCF in early childhood, alongside *Staphylococcus aureus*, although detection has been waning in recent years<sup>78</sup><br>- Long term colonization with hyper-mutable *H. influenza* strains has been<br>found to be more common in pwCF than the general population<sup>79</sup> |
| *Mycobacterium* | *Mycobacterium abscessus* | Bacterium | Lung | - Increasing in prevalence and associated with dramatic disease progression<sup>80</sup><br>- Often requires extensive antibiotic treatment<sup>80</sup><br>- Can be a contraindication for lung transplant<sup>80</sup> |
| *Pseudomonas* | *Pseudomonas aeruginosa* | Bacterium | Lung | - With increasing age, *P. aeruginosa* tends to dominate the airway, evolving to resist antibiotics and out-competing other airway pathogens<sup>70</sup><br>- Once chronic infection is established, constant antibiotic use is required to keep the bacterium at bay; once established it can rarely be eradicated<sup>81</sup><br>- *P. aeruginosa* infection has been associated with worsening lung function over time<sup>82</sup> |
| *Porphyromonas* | Multiple | Bacterium | Lung | - The genera *Porphyromonas* is commonly detected in the CF lungs<sup>83</sup><br>- Recent studies suggest that certain *Porphyromonas* species (namely *P. catoniae*) are abundant in pwCF who are not colonized with *P. aeruginosa*, and that significant decline in *P. catoniae* may serve as a biomarker for the onset of *P. aeruginosa* infection<sup>5</sup> |
| *Staphylococcus* | *Staphylococcus aureus* | Bacterium | Lung | - *S. aureus* often colonizes the lungs of pwCF at a relatively early age, but has been found to co-exist in some cases with *P. aeruginosa* in older people<sup>84</sup><br>- pwCF infected with both *S. aureus* and *P. aeruginosa* tend to exhibit worse lung function than those infected with either *P. aeruginosa* or *S. aureus* alone<sup>85</sup><br>- pwCF infected with *S. aureus* alone (vs. those with *P. aeruginosa* alone) tend to have better clinical outcomes<sup>85</sup> |
| *Stenotrophomonas* | *Stenotrophomonas maltophilia* | Bacterium | Lung | - *S. maltophilia* has a known capacity to persistently colonize the CF lung and to become multidrug resistant<sup>86,87</sup> |
| *Streptococcus* | Multiple | Bacterium | Lung | - In young children with CF, before the lungs are dominated by the classic CF pathogens (e.g., *P. aeruginosa*, *S. aureus*), *Streptococcus* species tend to be the predominate species<sup>88</sup><br>- The clinical implications of *Streptococcus* infection for pwCF are complex – infection with certain species like the *S. milleri* group are associated with pulmonary exacerbation (acute, rapid decline in lung function), yet enhanced relative abundance of *Streptococcus* species in the lungs is associated<br>with more mild lung disease<sup>89</sup> |

Decades of prior CF pathogen research has helped advise modern clinical treatments, and this published body of research continues to serve as a source of knowledge for drug development as well as inspiration for future studies. High-throughput transcriptomics experiments – of which RNA-Seq studies have recently become most common – are especially useful as a source of published data to inform future experiments [Figure 1a,b]. The global nature of transcriptomics data – i.e., the fact that it provides a snapshot of most/all genes at once – allows for the same gene to be compared across studies. In an ideal world, CF pathogen researchers would be able to view which microbial strains, treatment conditions, and media have previously been utilized, and perform a quick visualize analysis of gene expression under these conditions. This information would offer researchers a roadmap to identify future directions for follow-up experiments. However, we do not (yet) live in this ideal world. Although many data sets are publicly available, substantial computational expertise and manual effort are required to compare similar studies, visualize gene expression patterns within studies, and use published data to generate new experimental hypotheses. Thus, there is a need develop an application that will reduce these barriers to data re-analysis.

**Figure 1.**
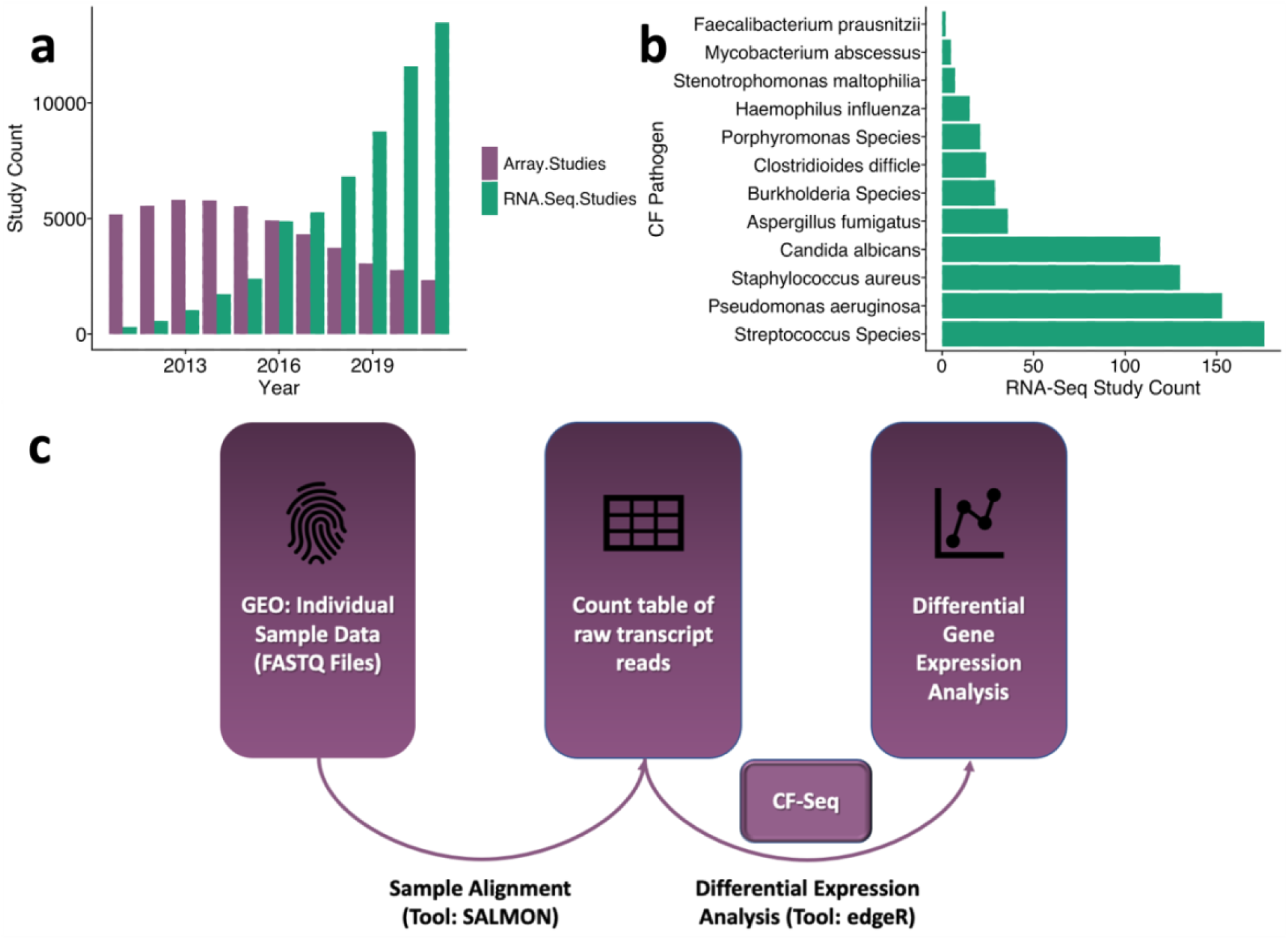
Landscape of RNA-Sequencing studies available in the Gene Expression Omnibus (GEO). (a) Since 2011, the number of RNA-Seq studies hosted in GEO has risen dramatically, from several hundred to over 10,000, well eclipsing the number of microarray expression studies currently produced per year. (b) While small relative to the total set of RNA-Seq studies in GEO, there is a substantial number of RNA-Seq studies available for the CF pathogen species featured in the CF-Seq application. (c) To derive meaningful biological insights from the RNA-seq studies in GEO, the analysis pipeline outlined here must be followed. Alignment of sample RNA sequences to a reference genome is accomplished with a command line tool like SALMON, and downstream analysis with a tool such as the popular R package edgeR. CF-Seq automates the second segment of this pipeline, saving users from the need to clean up count tables, produce experimental design matrices, gather metadata, and write sophisticated analysis code in R.

One useful approach to derive biological insights from a dataset in GEO – and the one that we automate in the CF-Seq application – is to see which genes are differentially expressed under varying experimental conditions. To accomplish this analysis, a researcher would first need to locate the sample runs associated with the individual dataset. These are often stored as FASTQ files that require extensive computational skills to process. Someone with these skills could trim the sequence reads contained in the FASTQ files to remove low quality reads and adapter sequences, and align trimmed reads to a reference genome with a command-line tool like SALMON^20^, which yields a count table with raw gene expression counts for each sample. Then, finally, that researcher could conduct differential gene expression analysis. This final step requires knowledge of a programming language like R^21,22^, and specific R packages like edgeR^23,24^ or DESeq^25^ that allow for the production of biologically meaningful analysis tables and figures. Even among bioinformatics researchers, many do not have expertise in all aspects of this pipeline – and for those who do, running through the pipeline for just a single data set is typically a multi-day effort. CF-Seq has been designed so that users do not have to deal with this pipeline at all. Taking advantage of count tables that dataset contributors have left in GEO as supplemental files, CF-Seq takes care of differential expression analysis [Figure 1c].

Our efforts to make public data more accessible are certainly not the first of their kind. In recent years, as big-omics datasets have become increasingly commonplace and researchers have encountered the challenges described above, the necessity of adopting FAIR data principles by making data sets more <u>F</u>indable, <u>A</u>ccessible, Interoperable, and <u>R</u>eproducible has increasingly been recognized^26^. In this spirit, various research tools have already been developed to make publicly available data more amenable to re-use. For example, the application *MetaRNA-Seq* enables users to view consolidated study metadata that had been scattered across the four NCBI databases: SRA, Biosample, Bioprojects, and GEO^27^. Another application, the *geoCancerPrognosticDatasets Retriever*, allows users to use additional search parameters (e.g., cancer type) to retrieve GEO accessions for all studies of interests^28^.

Some existing applications designed by other research teams are actually quite similar in nature to CF-Seq and have served as strong inspiration for our own efforts. However, none are specifically geared towards CF pathogen research, and there is room to expand on their functionality [Table 2]. Our own lab has previously published tools to make publicly available data more accessible to CF researchers^29,30^, but these tools focus on the most commonly studied CF pathogens – namely *Pseudomonas aeruginosa* and *Staphylococcus aureus* – and don’t include data sets on many of the other clinically relevant species listed in Table 1.

**Table 2.**
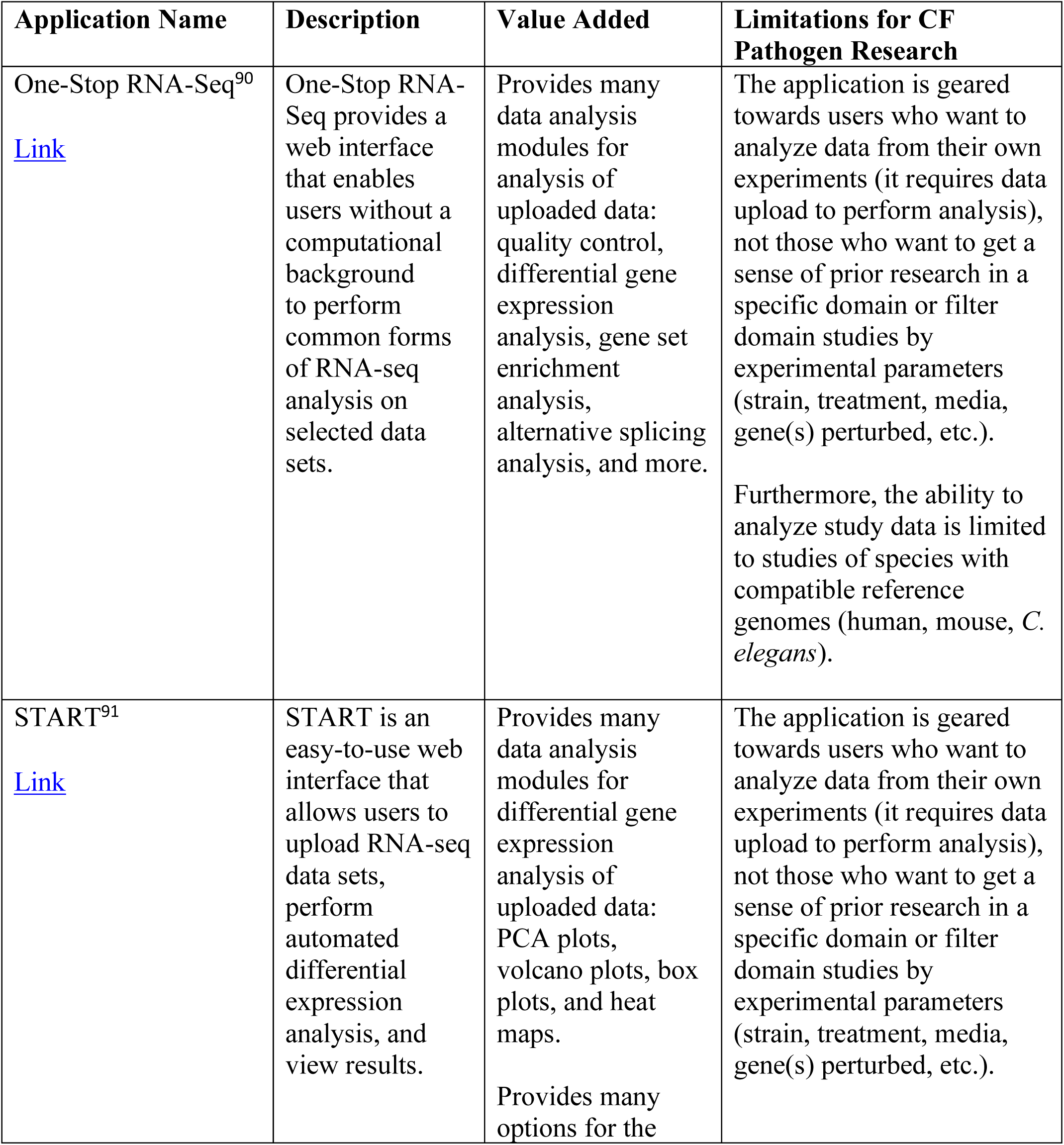
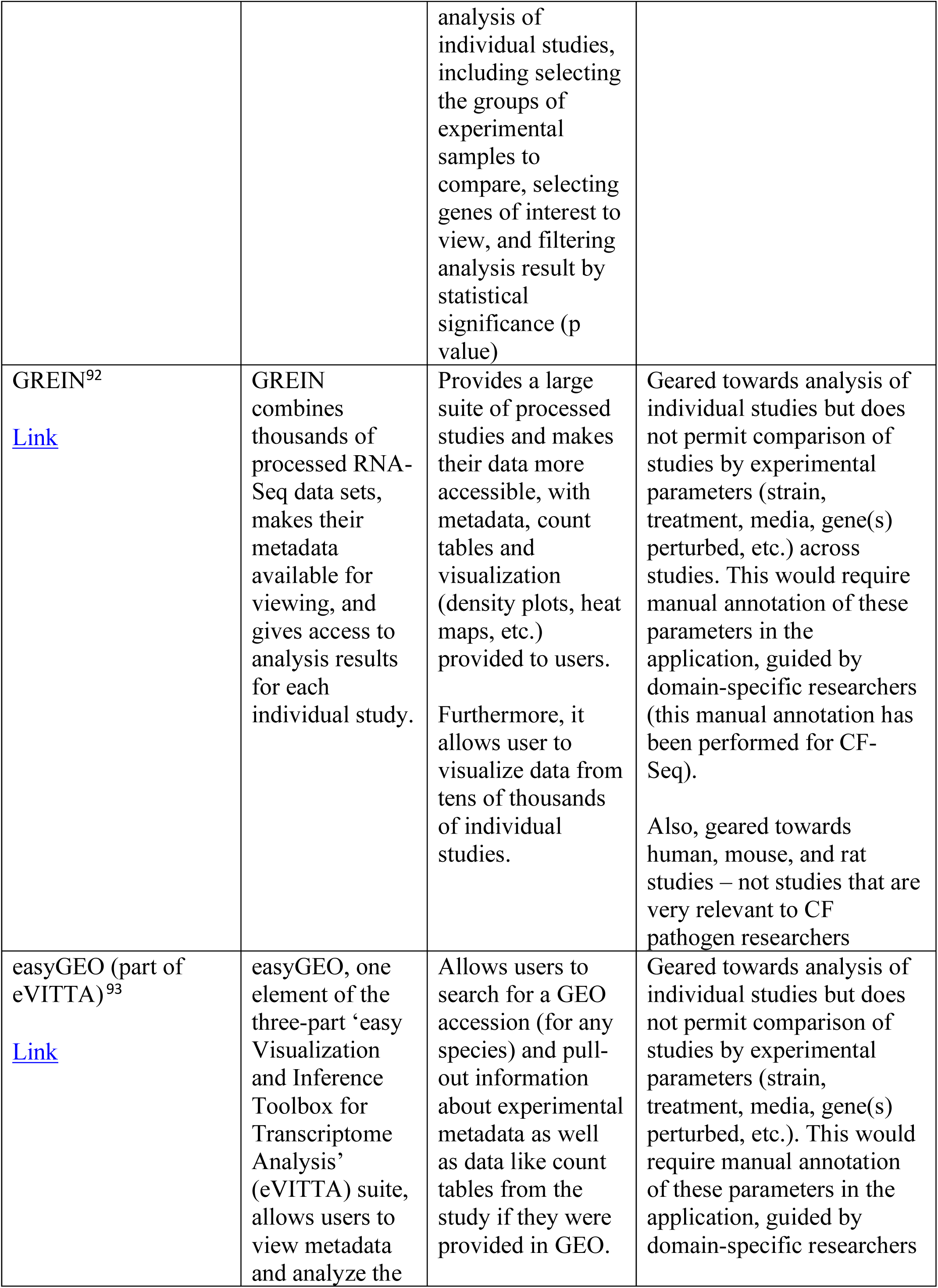
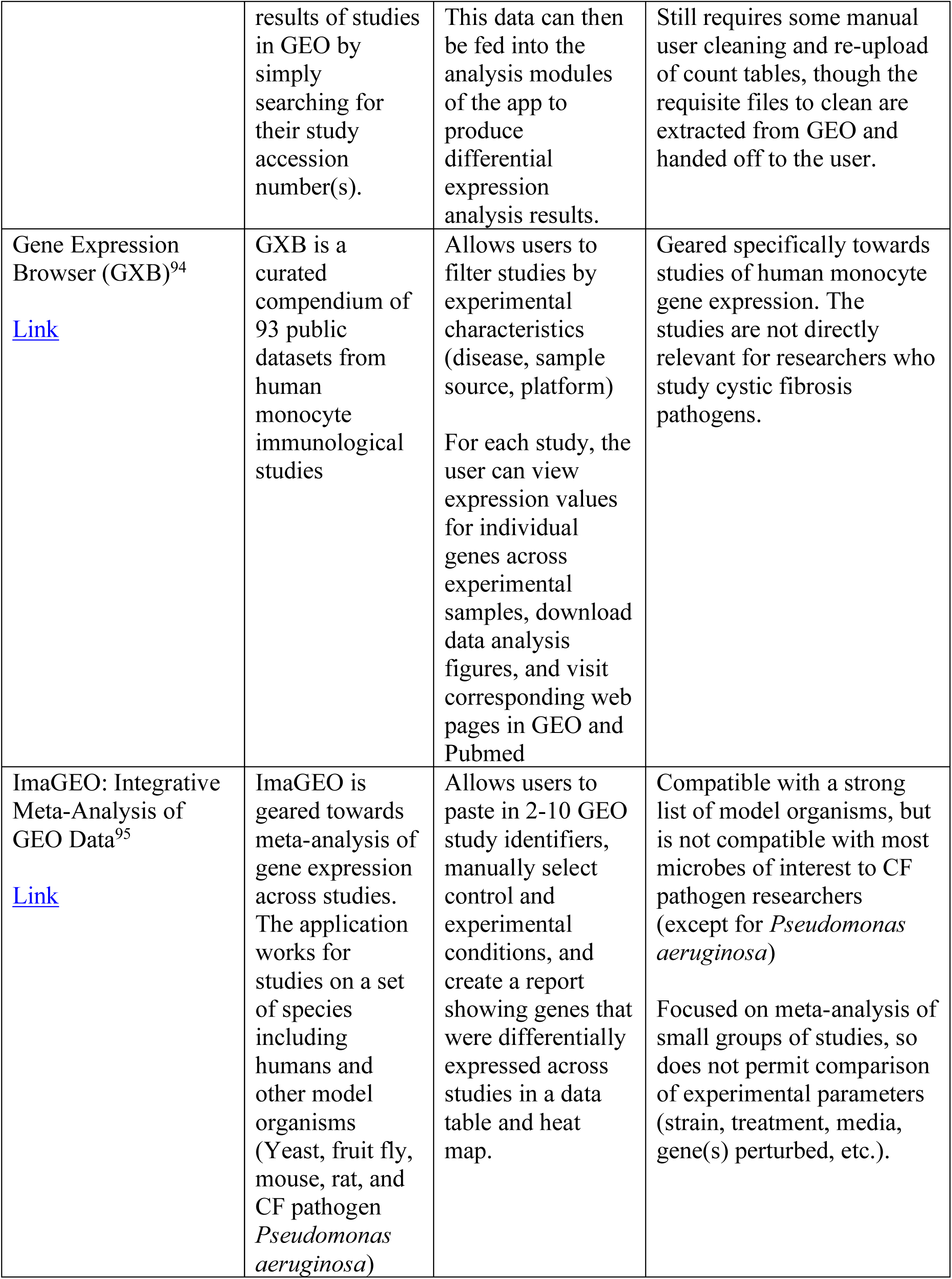
Applications similar in nature to CF-Seq are described to acknowledge the significant contribution that these researchers have made to making public data more FAIR, and the inspiration their work has provided for CF-Seq. The table also briefly summarizes how these applications are limited for CF pathogen research specifically to demonstrate the unique value of CF-Seq.

Building on our prior work, we present the R Shiny web application CF-Seq. CF-Seq is a web application based on a compendium of RNA-Seq experiments. This compendium contains 13 clinically relevant CF pathogens; a mix of aerobes and anaerobes residing in the lung and the digestive tract. The application currently holds carefully formatted count tables and metadata for 147 studies, and 1446 RNA-seq samples in total, with efforts ongoing (outlined in the Discussion section) to capture more studies and additional relevant species. All datasets currently included in the application are arranged by GEO accession number in supplemental table S1 for reference.

The CF-Seq application allows differential gene expression analysis of each individual study at the click of a button, producing downloadable tables and figures depicting fold changes and p values of differentially expressed genes in a matter of seconds. For each study, the application allows users to produce tables and figures comparing individual sample groups (e.g., samples treated with antibiotic X vs. control samples, samples treated with antibiotic Y vs. samples treated with antibiotic X, etc.). For many species and strains (where KEGG pathway annotations are available) the user can also visualize how the genes in specific biological pathways are differentially expressed. Furthermore, the user can filter all studies on the same species – breaking them down by strain, media, treatment, or gene(s) perturbed – to identify all past experimental conditions (and combinations of conditions) and thus determine which have yet to be assessed [Figure 2]. This application has been developed with the close guidance of CF pathogen researchers at the Geisel School of Medicine at Dartmouth College. In this publication, we present three case studies that showcase the application’s usefulness for researchers studying three different CF pathogens (*Aspergillus fumigatus, Pseudomonas aeruginosa*, and *Staphylococcus aureus*).

**Figure 2.**
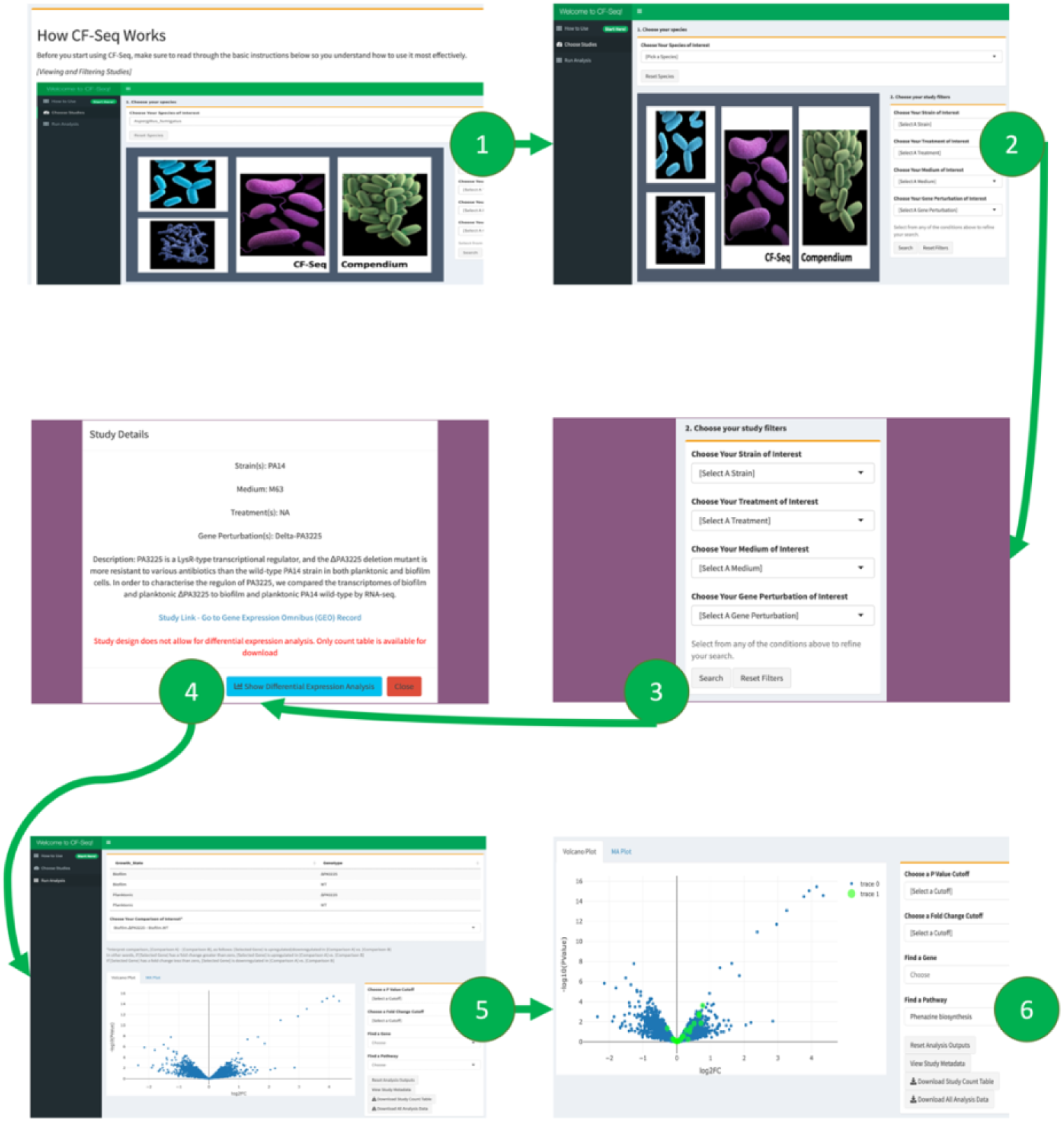
Application workflow for CF-Seq users. Panel 1 shows the starting window of the application, where users are presented with a manual that explains the functionality and purpose of the application. Users are then directed to the study view screen, shown in panel 2, where they can select a species of interest and view available RNA-Seq studies. Panel 3 shows how filters can be applied to delineate studies with certain experimental characteristics (strain, media, treatment, gene perturbed). Panel 4 offers a look at the metadata that can be examined for each individual study. Panels 5 and 6 show the study analysis window, where analysis tables and figures can be generated for all experimental comparisons, individual genes may be highlighted, P value and fold change cutoffs can be selected, and differentially expressed genes on selected KEGG pathways can be highlighted when KEGG pathway information is available (Panel 6). Zoomed-in versions of the figure panels showing more detail are available as supplementary figures S1-S6.

## Results

The CF-Seq application makes it simple for CF researchers to take full advantage of the 147 CF RNA-Seq data sets in the associated compendium. Upon opening the application, the user is greeted with a user manual that instructs them on how best to use CF-Seq (Figure 2, Panel 1). After reading, the user is then directed to the central, study-filtering panel of the application (Figure 2, Panel 2). Here, the user can filter studies by species, and then by strain, media, treatment, or gene perturbation (Figure 2, Panel 3). Filtered studies are presented in a table and can be selected to reveal additional metadata – including the study name, description, and link to its record in GEO (Figure 2, Panel 4). Once a study is selected, the user can click a button to reveal detailed differential expression analysis in a separate analysis tab (Figure 2, Panel 5). This analysis includes a table with the fold change (FC), p value, and counts per million (CPM) of all genes assessed in the study. For species or strains in which KEGG pathway information is available, the user is also able to visualize how the genes on different KEGG pathways are up or downregulated (Figure 2, Panel 6).

A series of user stories have been developed by three of the publication co-authors to demonstrate the value of the application in a research setting. These co-authors conduct research in laboratories that frequently publish papers related to CF microbiology. The following section of the manuscript demonstrates the analysis features of the application and outlines how these researchers used the application to come up with new questions and testable hypotheses relevant to their own research. Given the current focus in the field of CF research on the CF microbiome as a polymicrobial community^18,19^, all three user stories focus on polymicrobial interactions between several CF pathogens. All volcano plots used as figures for the user stories were taken directly from the application.

### Case Study #1: Examining *Aspergillus fumigatus* in bacterial co-culture

*Dr. Charles Puerner, Cramer Laboratory, Geisel School of Medicine*

The infectious mold *Aspergillus fumigatus* is ubiquitous in the environment^31^. The spores from this fungus are taken into the lung by breathing and normally cleared by a healthy immune system. However, individuals with compromised immune systems and pulmonary diseases such as cystic fibrosis are particularly vulnerable to infection by this fungus. In these cases, *A. fumigatus* spores are capable of germinating in the lung environment and forming fungal lesions. The Cramer lab studies the biology of this organism, specifically as it relates to its disease-causing capabilities. A recent publication, for example, investigated the genetic characteristics of persistent isolates taken from the lungs of a CF patient over several years^32^.

Using the analysis capabilities of this application, we were particularly interested in a dataset which generated gene expression profiles of *A. fumigatus* co-cultured with the ubiquitous bacterium *Pseudomonas aeruginosa* (GEO: GSE122391). This dataset is interesting because both organisms are commonly found in the CF lung environment, a situation associated with worsened disease state^33^. The study was identified using the CF-Seq filtering feature to focus on those experiments that involved cross-species interactions.

In the analysis window of the application, the “Choose a P Value Cutoff” field was used to highlight genes whose p-value was 0.05 or less. Genes were highlighted at several timepoints comparing the co-culture of *P. aeruginosa* with *A. fumigatus* to culture of the fungus alone (Figure 3). Volcano and MA plots demonstrating the magnitude of differential expression, as well as a spreadsheet of statistically significant differentially expressed genes, were quickly downloaded for further analysis and additional figure generation [Figure 3]. Then, the downloaded table of differentially expressed genes was easily filtered outside of the application to contain only genes with a |log_2_FC| value of 1.5 or greater (Fold change > 2.83 = log2FC > 1.5, Fold change < 0.35 = log2FC < −1.5) [Supplemental Table S2].

**Figure 3.**
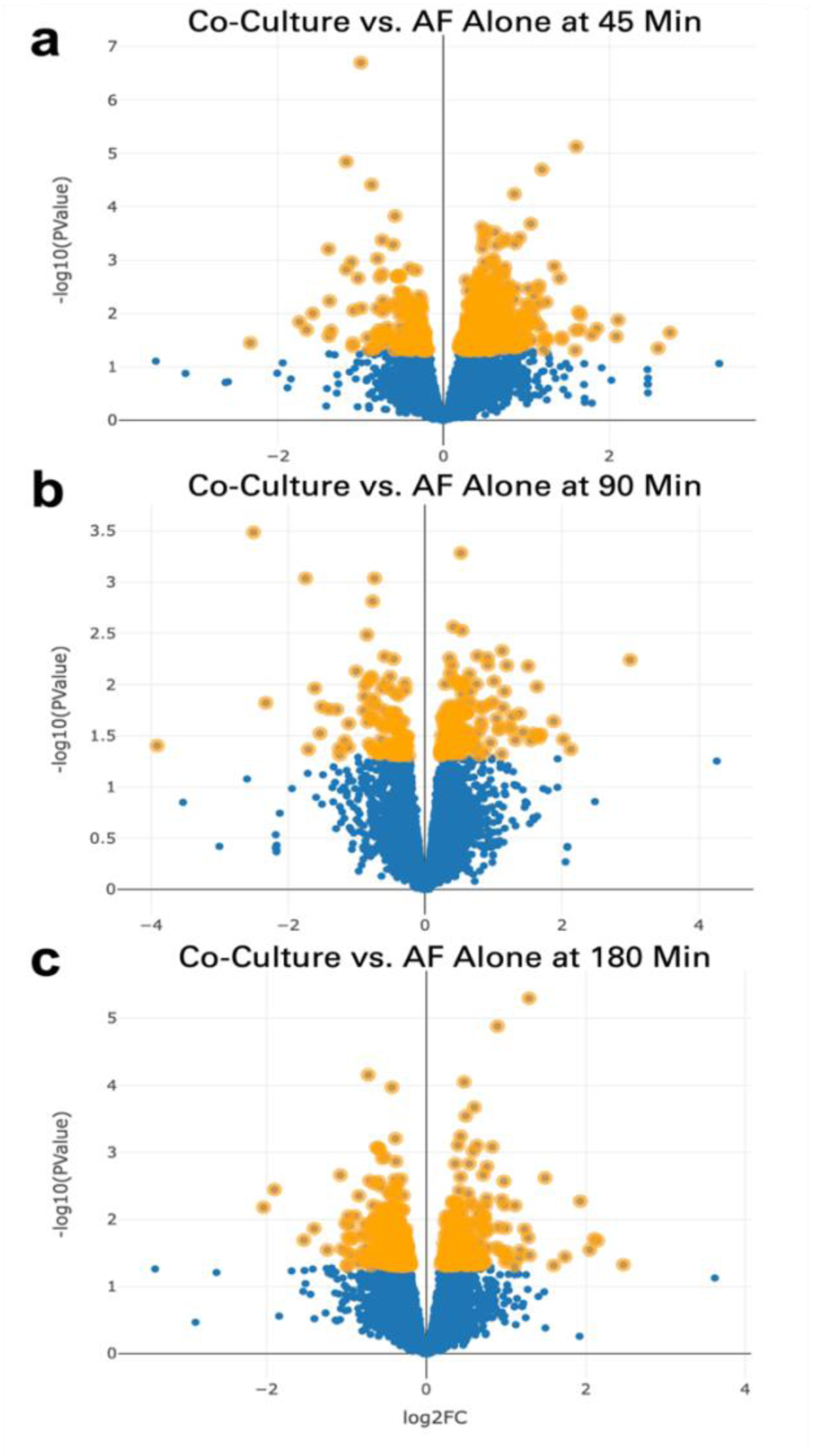
Expression of A. fumigatus genes following exposure to P. aeruginosa presented in volcano plot format. In the CF-Seq application, the species A. fumigatus strain A1160 was selected and the dataset “Transcriptomics analysis of Aspergillus fumigatus co-cultivated with Pseudomonas aeruginosa” was used for the subsequent analysis. Comparisons were selected comparing fungus co-cultured with bacteria to fungus alone at (a) 45, (b) 90, and (c) 180 min. Genes highlighted in orange are those whose p-value was less than 0.05. At 45 minutes, 531 of 8526 total genes were differentially expressed to a statistically significant degree. At 90 minutes and 180 minutes, the number of statistically significant differentially expressed genes was 257 and 514 respectively.

Using this method, the application makes it easy to identify a list of biologically significant genes which could be investigated further regarding their role in the co-culture environment (Figure 3). Genes differentially expressed with an especially high fold change and p value may be manipulated in the laboratory to see how the knockout of individual genes effects survival fitness of A. fumigatus in co-culture.

### Case Study #2: *P. aeruginosa* virulence factor production in polymicrobial contexts

Dr. Georgia Doing, Hogan Laboratory, Geisel School of Medicine

*Pseudomonas aeruginosa* is one of the most common pathogens associated with cystic fibrosis (CF) lung infections, remains difficult to treat with antibiotics, and is associated with lung function decline in colonized pwCF^34^. Along with its ability to form recalcitrant biofilms and resist antibiotic treatment, its behaviors during interactions with other bacteria are now recognized as important factors that influence *P. aeruginosa* infection outcomes^35,36,37,38,39^.

Microbial interactions are often studied in the laboratory using co-cultures of *P. aeruginosa* with other CF pathogens such as *Candida albicans* and *Staphylococcus aureus*. These co-culture experiments have proven to be useful for modeling polymicrobial interactions. However, it is increasingly apparent that the combinatorial effects of environmental factors and pairwise and community-wide microbial interactions make for complex systems with many changing variables and a large search space^39,40,41^. In addition to conducting new experiments in the laboratory, the re-analysis of individual data sets related to bacterial co-culture and meta-analysis of multiple datasets will likely spur new experimental hypotheses and help provide evidence for existing theories of polymicrobial interactions.

Using CF-Seq it was easy to compare two datasets from experiments where *P. aeruginosa* was co-cultured with *C. albicans* (GEO: GSE148597)^40^ and (GEO: GSE122048) *S. aureus*^42^. We noticed that while *P. aeruginosa* mainly upregulates and highly expresses genes in the KEGG pathway for phenazine biosynthesis in co-cultures with *C. albicans* compared to monoculture (Figure 4a), it does not do so in co-culture with *S. aureus* compared to monoculture (Figure 4b). Since *P. aeruginosa* phenazine production is induced with *C. albicans* fermentation^43^, we searched for specific genes whose expression could indicate differences in *C. albicans* and *S. aureus* metabolisms that may shed light on their different effects on *P. aeruginosa* phenazine production.

**Figure 4.**
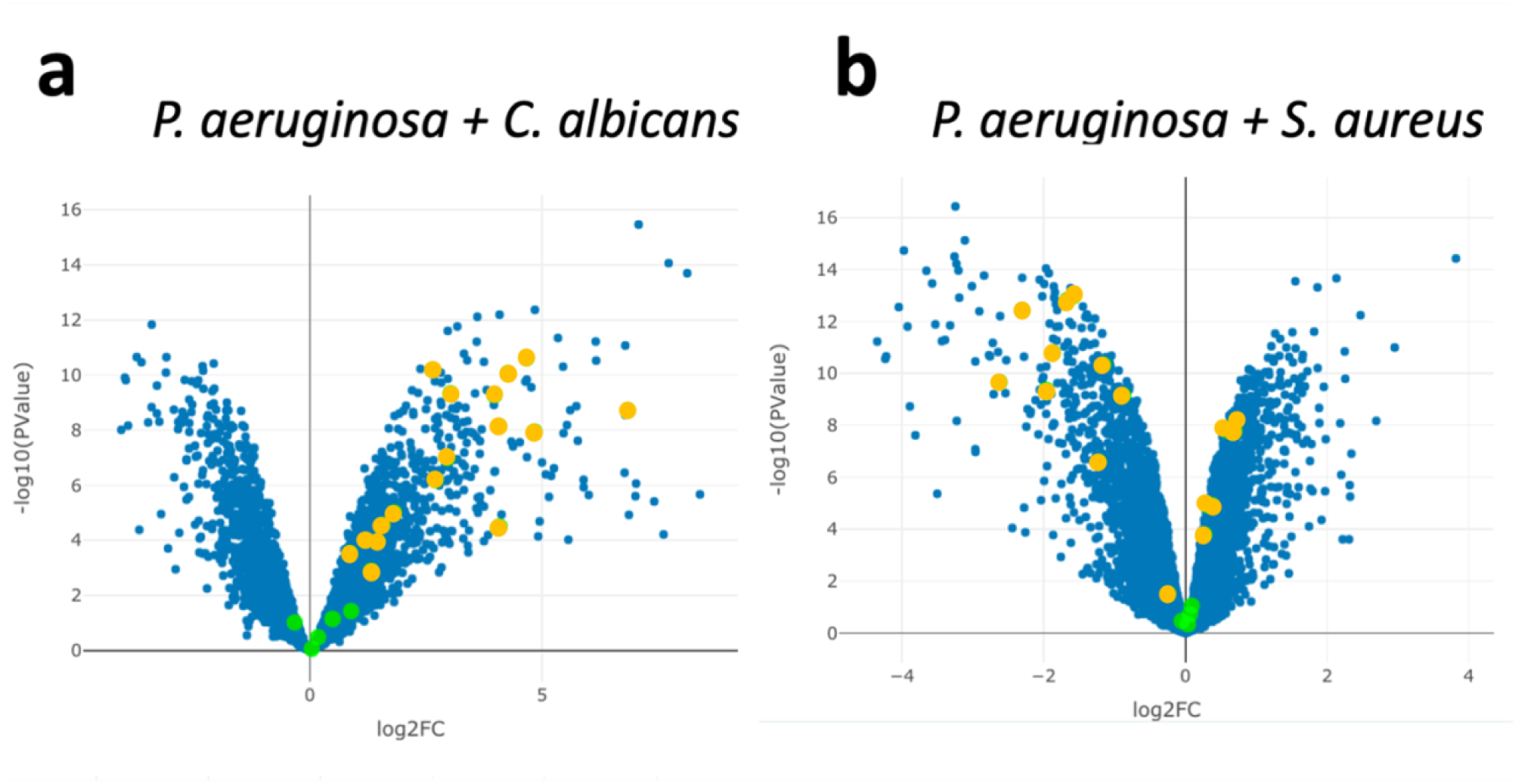
(a) P. aeruginosa genes involved in phenazine biosynthesis tend to be upregulated in co-culture with C. albicans (b) but not in co-culture with S. aureus compared to P. aeruginosa in monoculture. Green data points were highlighted by selecting the KEGG pathway for phenazine biosynthesis using the ‘find a pathway’ feature in the CF-Seq application. Genes that are differentially expressed between co-culture and monoculture conditions to a statistically significant degree (p < 0.05) were colored orange for emphasis.

Digging deeper into the data on an individual gene level, the upregulation of lactate permeases and lactate dehydrogenases by *P. aeruginosa* in co-culture with either *C. albicans* or *S. aureus* suggest both *C. albicans* and *S. aureus* were producing lactate in these experiments (Figure 5a). However, while *P. aeruginosa* upregulated alcohol dehydrogenases in co-culture with *C. albicans*, it did not do so in co-culture with *S. aureus*, suggesting *C. albicans* was likely producing ethanol while *S. aureus* was not (Figure 5b). Amongst the many differences between these two co-cultures, differences in microbially-produced fermentation products could lead to differences in *P. aeruginosa* phenazine production.

**Figure 5.**
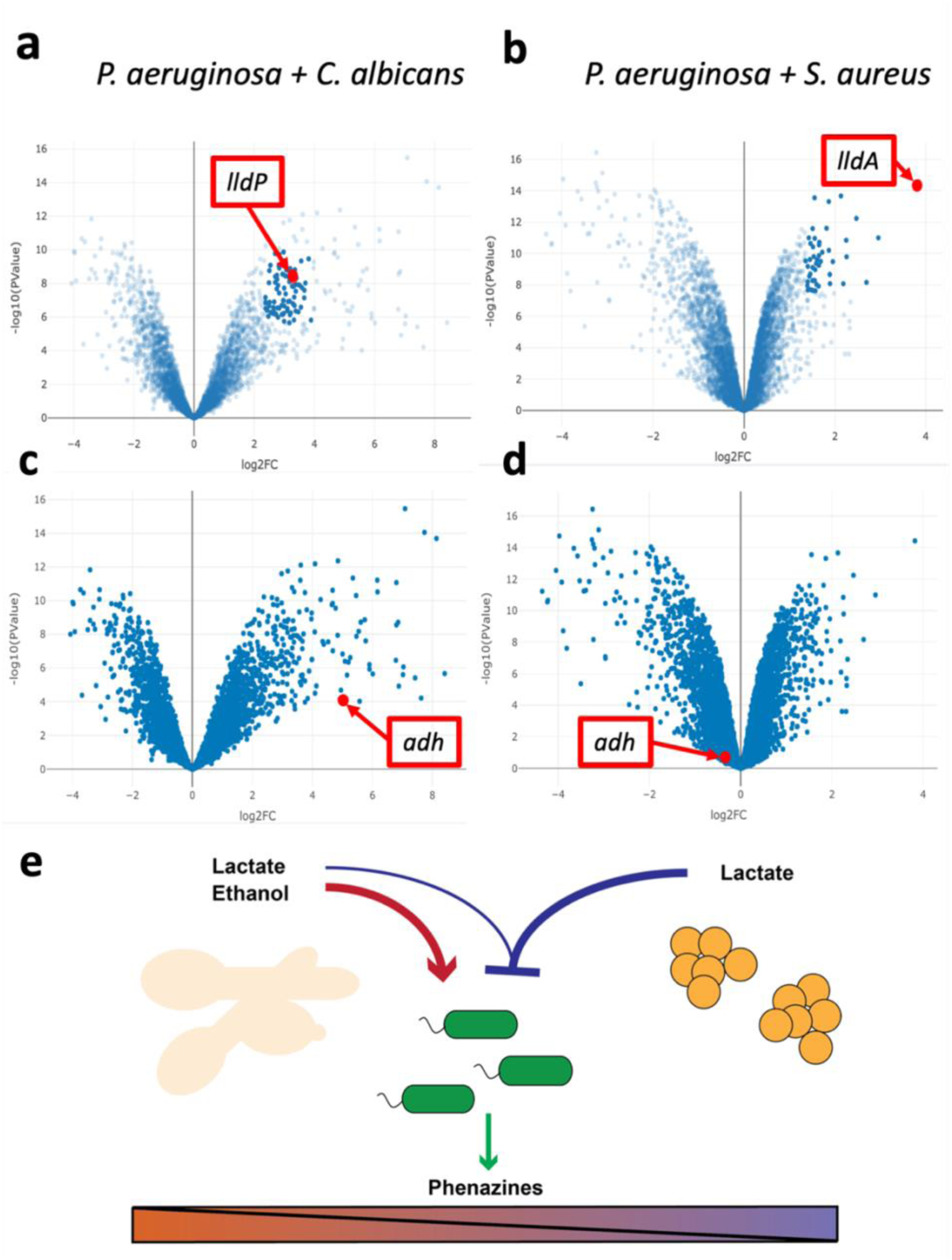
(a) P. aeruginosa upregulates the expression of lactate permease lldP (red point) and other lactate metabolism genes including lactate dehydrogenases (present in the cluster of dark blue points near lldP) in co-culture with C. albicans. (b) Similarly, lactate dehydrogenase lldA (red point) and other lactate metabolism genes (included in dark blue points near lldA) are upregulated in co-culture with S. aureus as well. (c) P. aeruginosa upregulated alcohol dehydrogenase adh in co-culture with C. albicans (d) but not in co-culture with S. aureus. (e) In complex co-culture P. aeruginosa will have to integrate multiple signals such as the positive influence of ethanol and a possible negative influence of lactate that converge to influence phenazine production. After CF-Seq exploratory analysis, our hypothesis is that the presence of ethanol will supersede that of lactate to promote phenazine production.

Since both co-cultures elicited lactate metabolism, but only co-culture with *C. albicans* elicited ethanol metabolism, CF-Seq analysis suggests that ethanol specifically promotes phenazine production while lactate may have a neutral or repressive effect (Figure 5c). This hypothesis could easily be tested in the lab by the addition of sub-lethal concentrations of ethanol to *P. aeruginosa* and *S. aureus* co-culture and measuring phenazine biosynthesis to test the hypothesis that phenazine production would increase. Importantly, CF-Seq facilitated the re-analysis of public data that led to the development of a hypothesis in approximately 30 minutes. By contrast, the process of identifying these experiments, downloading the data, performing comparisons, and generating figures by hand would have taken approximately 16 hours, based on similar exploratory analyses that we have performed previously.

### Case Study #3: Examining superoxide dismutase response in *Staphylococcus aureus* under a variety of clinically relevant conditions

Liviu Cengher, Cheung Laboratory, Geisel School of Medicine

*Staphylococcus aureus* is a human commensal and opportunistic pathogen that contributes to a wide range of diseases – from skin and soft-tissue disorders to respiratory diseases like cystic fibrosis^44^. Disease is mediated by several *S. aureus* virulence factors that are produced in response to environmental cues, and which play a wide range of roles^45^. Two-component systems (TCS) are important regulatory factors that have paired sensing and regulatory peptides that respond to environmental and host cues^46,47^. The *SaeR/S* TCS senses reactive oxygen species (ROS) and regulates responses that counteract and inhibit ROS production by the human immune system. For example, activation of the TCS may lead to enhanced expression of virulence factors superoxide dismutase *sodA* and *sodM*^48^.

In this case study we investigated *sodA* and *sodM* expression across experiments with different bacterial strains and treatments to explore similarities and differences in ROS response. Specifically, we compared *sodA* and *sodM* expression in conditions likely to be present in the CF lung to identify conditions that upregulate one and/or both of the two genes. To start, we evaluated the effect of *S. aureus* co-culture with *P. aeruginosa* (vs. *S. aureus* in monoculture, GEO: GSE122048)^42^. Co-occurrence of *P. aeruginosa* and *S. aureus* is frequent in a hospital setting, and tends to induce a fermentative state in *S*.*aureus*^49,50^. Both *sodA* and *sodM* were upregulated in these conditions (Figure 6a,b). CF-Seq analysis of the transcriptome of ‘persister cells’ primed to survive (predominantly ROS mediated) killing after residing inside of immune system macrophages (GEO: GSE139659)^48,51^ revealed that *sodM* was upregulated in the ‘persister cells’ that resisted killing by the immune system (Figure 6d). Since *sodM* was upregulated in common between these two studies, it would be worth re-examining both conditions in tandem: subjecting *S. aureus* to bacterial co-culture with *P. aeruginosa* to see if this induces a persister-like phenotype in *S. aureus*. Given that both conditions – persistence within host cells and co-infection with *Pseudomonas* – may be present at once in an individual with CF, such experiments would paint a fuller picture of the *S. aureus* transcriptional state in an infection.

**Figure 6.**
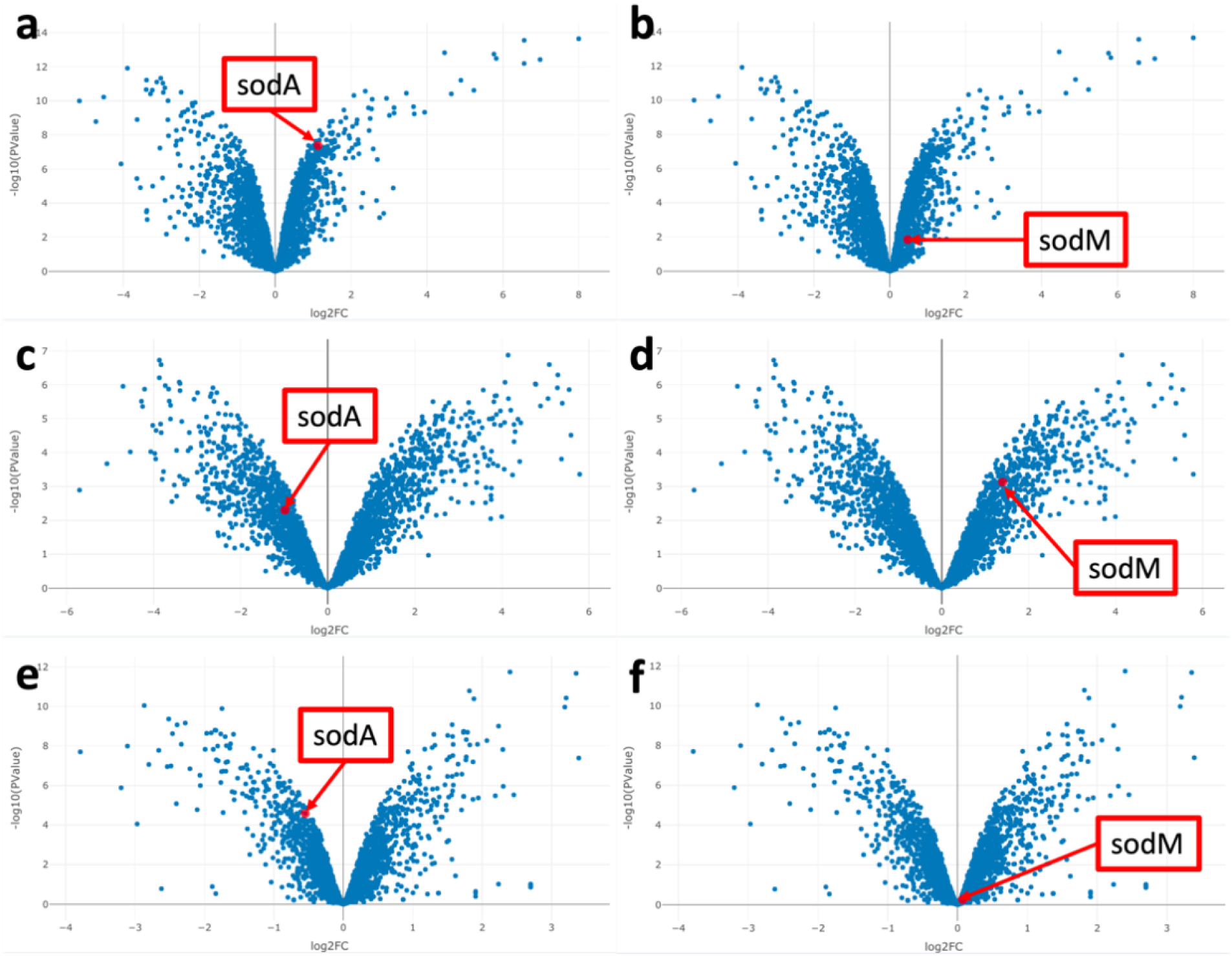
Expression of virulence factors sodA and sodM in S. aureus tends to diverge under different experimental conditions. Volcano plots of all genes are shown to demonstrate the expression values of sodA and sodM relative to other genes detected. (a,b) In co-culture with P. aeruginosa, both sodA and SodM expression are upregulated, sodA to a much greater extent. (c,d). In ‘persister cells’, the expression pattern was quite different: sodM expression was more markedly upregulated while sodA expression was downregulated (e,f). Finally, exposure to apicidin was found to induce downregulation of sodA, but no significant change in sodM. In all cases, aside from sodM expression in figure 6f, sodA and sodM were differentially expressed to a statistically significant degree (p < 0.05)

Furthermore, we also identified a study where treatment with apicidin, an antibiotic known to inhibit bacterial quorum sensing, led to downregulation of *sodA* and relatively low levels of *sodM* expression (Figure 6e,f)^52^. We might compare *sodA* downregulation in this study with the co-culture study (GEO: GSE122048). There would be interesting therapeutic implications for future experiments that determine the outcome of combining co-culture conditions (upregulating *Sod* genes) with antibiotic-induced quorum sensing inhibition (downregulation/low expression of *Sod* genes) to see which effect dominates. In addition, one might examine conditions which could favor *sodM* expression over *sodA* expression, like the availability of iron and manganese in co-culture and in polymicrobial infections^53,54^. Normally the analysis performed in this case study would necessitate a close reading of multiple published articles and require deciphering often unhelpful supplemental data tables. Finding relevant experiments and performing subsequent analysis would involve many hours of work. Using CF-Seq, useful results were found within approximately 10 minutes.

## Discussion

As the user stories demonstrate, the CF-Seq application provides value to CF pathogen researchers in a number of ways. First, CF-Seq allows rapid analysis of numerous data sets, reducing the time of analysis from days in some cases to minutes. Second, multiple CF pathogens can be analyzed including bacteria such as *Pseudomonas aeruginosa* and *Staphylococcus aureus*, and fungi such as *Aspergillus fumigatus* and *Candida albicans*. Third, the R scripts underlying the application (publicly available in our Git Repository: https://github.com/samlo777/cf-seq.git) not only allow for rapid analysis of the CF pathogens currently included in the application, but may be repurposed to study other microbes relevant to other diseases. Fourth, CF-Seq affords researchers a better understanding of prior CF pathogen experiments by revealing experimental parameters – details on strain, media, treatment, and gene perturbation – that have been tested in the past. With the ability to filter studies based on these parameters, users may identify the set of experiments that relate to their own specific interests and capabilities, filling knowledge gaps that they notice in the field of research. Not only does the application make prior studies more visible and accessible, but also makes their individual samples and the expression levels of individual genes possible to investigate more closely. While any given publication tends to emphasize the differential expression of just a few relevant genes to tell a concise and cohesive biological story, the CF-Seq application allows users to explore the expression of genes that may not have been of interest to the initial study authors but are of interest to the users themselves.

The ability to discern the whole field of prior experiments in minutes without slowly trawling through online databases like GEO is a tantalizing prospect. As it stands, the application serves as a valuable tool for validating existing hypotheses and generating new ones to test. That said, efforts are still ongoing to expand the application – adding older microarray studies to the compendium of data and making efforts to gather count table data for RNA-Seq studies in which count tables have not yet been provided directly by the authors as supplemental information in GEO. Additional RNA-Seq studies may be gathered by taking advantage of pipelines built to convert FASTQ sample files in GEO into count tables amenable to analysis by edgeR. For example, we may employ the pipeline recently developed by Doing et al. (2022) to create a compendium of *P. aeruginosa* data sets, modifying it such that its use extends to other CF pathogens of interest^55^. We may also take advantage of crowd-sourced metadata curation approaches like that of Wang et al. (2016), in which participants were recruited to help identify studies in GEO involving gene or drug perturbations, or comparison of normal and diseased tissue^56^. Crowdsourcing curation efforts would make the process of adding additional study data to the application more efficient and speed up the inclusion of new studies.

Finally, the application sheds light on the value of automated bioinformatic analysis for researchers of all backgrounds. Performing differential expression analysis is by no means a feasible task for those lacking a computational background, and even for those who have such a background, analysis is still quite time-consuming (as the authors of the user stories note). Not only does the CF-Seq application save time and provide detailed statistical analysis, but it also serves a didactic purpose for those who have less experience working with transcriptomic data – demonstrating what differential expression analysis looks like and how it may be interpreted.

Tools such as CF-Seq, and the other data re-analysis applications cited throughout this publication, demonstrate the immense value of bioinformatic tools for scientific research. In sum, providing CF pathogen researchers a more detailed view of the prior experiments conducted in their own domain will make research more coordinated, systematic, and efficient. The CF-Seq application allows users to see exactly what combinations of experimental factors have been assessed thus far, and take logical, incremental steps – investigate a new treatment, a new mutation, a new growth medium, or some combination thereof – to test novel experimental hypotheses and improve understanding of pathogen behavior. For the field of bioinformatics specifically, such an application helps demonstrate the value and enhance appreciation for both data re-analysis and the tools that enable it. More generally, applications like CF-Seq help democratize the research process, allowing all scientists, regardless of specialization, to set their minds at work determining where research should go next.

## Methods

### Data Extraction

The CF-Seq application currently includes 147 RNA-Seq studies of 13 CF pathogens. All studies can be found in NCBI’s Gene Expression Omnibus (GEO) and are linked directly to GEO within the application. Before incorporating studies into the application, the landscape of CF pathogen studies in GEO was surveyed. Clinically relevant pathogens of interest were chosen based on the cystic fibrosis literature (their relevance, supported by clinical and laboratory studies, is documented in Table 1). The set of all RNA-sequencing studies for each of these pathogens was identified in GEO by querying the database of GEO data sets by pathogen name (e.g., *Pseudomonas aeruginosa, Staphylococcus aureus*, etc.), filtering studies to include only those that constituted “expression profiling by high throughput sequencing” (in GEO, this corresponds to ‘RNA sequencing’), and selecting the pathogen of interest specifically in the ‘organism’ field. This final step excludes datasets that constitute transcriptomic profiles of human cells, or cells of some other organism, exposed to the pathogen of interest.

For practical reasons, only studies with certain attributes are included in this release of the compendium. The application is limited to studies where: A) a count table was provided in the supplemental files associated with the study in GEO, B) that count table was in a tabular format (.csv, .xlsx, .txt) so that it could be loaded into R with the read.table() or read.csv() functions, C) sample groups were clearly distinguishable such that it was possible to perform differential expression analysis, and D) the count table included raw counts and not normalized counts (edgeR and other differential expression analysis packages require raw counts to perform analysis). Efforts to circumvent some of these limitations and add more studies into the application are discussed in the Discussion section of this manuscript.

### Data Cleaning and Storage

For studies that met the criteria for inclusion in the application, each count table was subjected to the following formatting protocol. Count tables downloaded directly from GEO were re-structured, if necessary, so that the first column of the table included gene names, and all subsequent columns contained raw read data for each experimental sample. In addition to count tables, two other data files were constructed for each study. The first file is a design matrix which delineates experimental samples by condition (e.g., control, treatment group X, treatment group Y) and lists the number of replicates for each condition. This design matrix is a requirement for differential expression analysis with edgeR. The second file, labeled ‘additional metadata’, includes manually gathered metadata on the strain(s), media, treatment conditions, and genes perturbed in each study, whenever applicable. Collecting this data enables filtering of studies by experimental conditions within the application.

All data files – count tables, design matrices, and additional metadata – were deposited in a local directory of folders, with a single folder for each species, and sub-folders within each species for the three types of data files (count table, design matrix, additional metadata). A copy of this directory structure can be found in the Git Repository associated with this publication [https://github.com/samlo777/cf-seq.git], so that any reader may download the data and/or use it to run the Shiny application on their own computer if they so choose.

### Code Development Approach

The CF-Seq application code was developed in discrete modules to make testing as straight-forward as possible. Each of the application’s interactive features (filtering studies, selecting a study, choosing experimental comparisons to analyze, etc.) were developed in a hierarchical fashion: the code was first tested to ensure that it worked properly for a single study, then adjusted and generalized such that it worked for a single species, and ultimately for all species included in the application.

The application code is broken down into 3 files. The first, named ‘app.R’, contains the functional code for the application. This file houses the UI code (which dictates the appearance of the application), and the server code (which provides functional code for all the drop-down menus, tables, and output figures) as two separate blocks. Both the name of this file, and the two-section structure, are an essential requirement of all Shiny applications. In addition, another code file, labeled ‘Data Setup.R’, was generated to load in all the study data and compress it into the easily accessible structure (a list of lists) accessible to the code in the app.R file. In addition to loading in the count tables, design matrices, and additional metadata, this code file also contains blocks of code that perform differential expression analysis – and deposit the outputs of this analysis (including tables of fold changes, p values, and counts per million for each gene) into the list of lists object alongside their respective studies. The third code file, labeled ‘PathwayData.R’, contains code that programmatically accesses data from the KEGG database and structures that data such that the genes of each species are linked to their respective KEGG biological pathway identifiers (if available). The data structures generated by this code file are necessary for the biological pathway feature of the application.

The application takes advantage of several publicly available, open-source R packages. Alongside the ‘shiny’ package^57^ (which is essential for all R Shiny applications), the ‘shinydashboard’^58^ package was used to provide a UI template, with several tabs for different application components. ‘shinyjs’^59^ was used to develop some of the more complicated application features (e.g., data tables with interactive buttons) that require JavaScript code to run. The ‘DT’^60^ package was employed to create searchable and filterable tables. The package ‘plotly’^61^ was used to generate interactive volcano plots and MA plots to represent differential expression analysis results, and the differential gene expression analysis itself was performed with the ‘edgeR’^23,24^ package. Finally, the ‘tidyverse’^62^ suite of packages, including ‘stringr’^63^ for string manipulation, were used throughout the application code to manipulate data structures.

### Validation: Beta Testing Protocol

To ensure that the study data and metadata loaded into the application recapitulated the data present in GEO, and that all application features worked as expected, a beta testing protocol was established. Three of the paper co-authors, each possessing either domain knowledge in CF microbiology or bioinformatics, were recruited to test different segments of the application: (1) the ability to filter studies based on experimental characteristics, (2) the ability to view detailed metadata for each individual study, and (3) the ability to perform and visualize differential expression analysis. The beta testing protocol was guided by a series of requirements tables that listed out all the features to be validated (beta testers were instructed to indicate Y/N if a feature worked as expected and provide notes if it did not). These tables are included for reference in the supplemental material [Supplemental Tables S3 – S5]

After all components of the application were tested, any features that did not work properly were fixed – and additional improvements were made to enhance the usability of the application based on beta tester feedback. Furthermore, after the bugs identified in beta testing were fixed, a second round of review was undertaken to ensure that study metadata accurately reflected the true study metadata in GEO. One at a time, each study in the application was referenced back to GEO to ensure that none of the manually curated metadata was missing or incorrect.

## Project Documentation

Documentation for the CF-Seq application can be found in several locations. Users are presented with a user manual when they first open the application. This guide can also be found as a .pdf file in the Git repository [https://github.com/samlo777/cf-seq.git], which also contains the application’s source code and a README file that outlines the repository contents.

## Data Availability

All data – including count tables derived from GEO, and manufactured design matrices and metadata tables – are available in the Git repository [https://github.com/samlo777/cf-seq.git]

## Code Availability

All CF-Seq code is open source and has been made available for use on GitHub [https://github.com/samlo777/cf-seq.git].

The application is also hosted on a server maintained by Dartmouth College and is accessible at the following web link [http://scangeo.dartmouth.edu/CFSeq/)].

In its current version, CF-Seq utilizes the following R package versions: *shiny* (1.6.0), *shinydashboard* (0.7.1), *shinyjs* (2.0.0), *DT* (0.19.1), *plotly* (4.9.4.1), *ggplot2* (3.3.5), *edgeR* (3.34.1), *tidyverse* (1.3.1), *stringr* (1.4.0).

## Acknowledgements

Support for these studies was provided by the Cystic Fibrosis Foundation (grants STANTO19R0 – CFF and CRAMER19GO – CFF), the NIH (grants P30 DK117469, R01 HL151385, and R01AI146121), and The Flatley Foundation.

## Author Information

### Contributions

S.N. wrote the publication except for the user stories, and conceived of, developed, and tested the CF-Seq application. T.H. provided valuable guidance and inspiration through the entire app development and publication writing process. K.K. provided helpful guidance on app development and handled hosting of the app on the online server. L.C., C.P., and A.L. all helped with beta testing the application. L.C., C.P., and G.D. provided user stories demonstrating how the application could be used to test wet bench hypotheses. B.S. provided domain expertise (specifically, on P. aeruginosa), guidance on publication, and primary funding for the project. A.C., D.H., and R.C. also provided domain expertise and guidance. All authors reviewed drafts of the publication and provided feedback.

