## Supplementary Tables and Figures for "CF-Seq, An Accessible Web Application for Rapid Re-Analysis of Cystic Fibrosis Pathogen RNA Sequencing Studies"

*Supplementary Information*

---

Contents

Table S1. Study Data Acknowledgements
Table S2. Example Table of Differentially Expressed Genes
Table S3. Beta Testing Criteria – Filtering Studies
Table S4. Beta Testing Criteria – Viewing Study Metadata
Table S5. Beta Testing Criteria – Running Analysis

Figure S1. Zoomed-in image of figure 2 (application workflow), panel 1 (user manual)
Figure S2. Zoomed-in image of figure 2 (application workflow), panel 2 (study view)
Figure S3. Zoomed-in image of figure 2 (application workflow), panel 3 (filter studies)
Figure S4. Zoomed-in image of figure 2 (application workflow), panel 4 (additional metadata)
Figure S5. Zoomed-in image of figure 2 (application workflow), panel 5 (analysis view)
Figure S6. Zoomed-in image of figure 2 (application workflow), panel 6 (visualize pathway)

---

**Table S1. Study Data Acknowledgements.** All published data featured in the CF-Seq application is listed in this table, alongside the data contributors cited in GEO and a link to the associated publication, if one exists.

| GSE ID | Species | Title | Contributor(s) | Associated Publication |
|---|---|---|---|---|
| GSE122391 | *A. fumigatus* | Transcriptomics analysis of *Aspergillus fumigatus* co-cultivated with *Pseudomonas aeruginosa* | Keller S, Heinekamp T, Valiante V, Brakhage AA | NA |
| GSE140840 | *A. fumigatus* | Sensitivity of transcription factor mutants of *Aspergillus fumigatus* to Congo Red | Raj S, Sismeiro O, Legendre R, Varet H, Bromley M, Latgé J | PMID: 34006660 |
| GSE152682 | *A. fumigatus* | Transcriptome analysis of conidium germination of *Aspergillus fumigatus* in different growth conditions | Danion F, Legendre R, Sismeiro O, Varet H, Mouyna I | PMID: 33419224 |
| GSE173349 | *A. fumigatus* | The peroxiredoxin Asp f3 acts as redox sensor in *Aspergillus fumigatus* | Boysen JM, Saeed N, Wolf T, Panagiotou G, Hillmann F | PMID: 33946853 |
| GSE55648 | *A. fumigatus* | Transcriptome study of *A. fumigatus* hyphae and human neutrophils | Lapp K, Vödisch M, Kroll K, Pähtz V, Bruns S, Strassburger M, Linde J, Guthke R, Kniemeyer O, Valiante V, Heinekamp T, Brakhage AA | NA |
| GSE55943 | *A. fumigatus* | Next Generation Sequencing identifies O2-responsive genes in the human pathogenic fungus *Aspergillus fumigatus* | Hillmann F, Linde J, Brakhage AA | PMID: 24948085 |
| GSE95798 | *A. fumigatus* | Transcriptome analysis of *Aspergillus fumigatus* in the presence of sugarcane bagasse | Gouvêa PF, Bernardi AV, Gerolamo LE, Santos Ed, Riano-Pachon DM, | PMID: 29614953 |

| | | | | |
|---|---|---|---|---|
| | | | Uyemura SA, Dinamarco TM | |
| GSE97495 | *A. fumigatus* | *Aspergillus fumigatus* protein phosphatase PpzA is involved in iron assimilation, secondary metabolite production, and virulence | Oliveira Manfiolli A, Alves de Castro P, Dolan S, Doyle S, Riaño Pachón DM, Ulaş M, Noble LM, Mattern D, Valiante V, Bayram O, Goldman GH | PMID: 28753224 |
| GSE97539 | *A. fumigatus* | Gene expression changes in the absence of the carbon catabolite repressor, CreA. | Beattie S, Cramer R | PMID: 28423062 |
| GSE129572 | *Bacteroides* species | Analysis of the transcriptome of *Bacteroides thetaiotaomicron* during growth on complex N-glycan as the sole carbon source | Bolam DN, Briliute J | PMID: 31160824 |
| GSE74379 | *Bacteroides* species | RNA-seq analysis of *Bacteroides xylanisolvens* XB1AT grown on pectin or xylan | White BA, Fields CJ, Mosoni P, Despres J | PMID: 27142817 |
| GSE147369 | *Burkholderia* species | Transcriptional of *Burkholderia thailandensis* E264 and *Burkholderia thailandensis* BPM | LI J, Zhong Q, Lu W | NA |
| GSE155982 | *Burkholderia* species | *Burkholderia cenocepacia* K56-2 transcriptomic profile during the early contacts with surface of host bronchial epithelial cells | Fialho AM, Pimenta AI, Bernardes N, Mil-Homens D | PMID: 33707642 |
| GSE19115 | *Burkholderia* species | Mapping the *Burkholderia cenocepacia* niche response via high-throughput sequencing | Yoder-Himes DR, Chain PS, Zhu Y, Wurtzel O, Rubin EM, Tiedje JM, Sorek R | PMID: 19234113 |
| GSE77970 | *Burkholderia* species | Comparison of the transcriptomes of a *Burkholeria* | Boyce JD, Powell D | PMID: 27147217 |

| | | | | |
|---|---|---|---|---|
| | | *pseudomallei* bprR [bpss0688] mutant, a bprS [bpss0687] mutant and a bprRS double mutant with the wild-type *B. pseudomallei* K96243 parent strain. | | |
| GSE100737 | *C. albicans* | Transcript levels of genes in the wild type CAI4 as well as the gdt1/gdt1,the pmr1/pmr1 and the gdt1/gdt1 pmr1/pmr1 mutants | Jiang L, Yang Y | PMID: 29954393 |
| GSE110291 | *C. albicans* | Characterization of a *Candida albicans* Isolated from a recurrent Cervical Lymphadenitis Patient and it's Clinic Indication | Zhang C | NA |
| GSE116533 | *C. albicans* | Interaction of soil amoeba *Protostelium aurantium* with three pathogenic yeasts *Candida* spp. (here: *C. albicans* and *C. glabrata*) | Radosa S, Linde J, Hillmann F | NA |
| GSE123412 | *C. albicans* | Gcn5 Controls Virulence of the Human Fungal Pathogen *Candida albicans* through Multiple Regulatory Networks | Shivarathri R | PMID: 31263212 |
| GSE136116 | *C. albicans* | Global transcriptomic analyses of the *Candida albicans* response to treatment with a novel inhibitor of filamentation | Romo JA, Zhang H, Hong C, Wang Y, Lopez-Ribot JL | PMID: 31511371 |
| GSE136940 | *C. albicans* | Humans and other commonly used model organisms are resistant to cycloheximide- | Sharma P, Nilges BS, Wu J, Leidel SA | PMID: 34429433 |

| | | | | |
|---|---|---|---|---|
| | | mediated biases in ribosome profiling experiments | | |
| GSE147697 | *C. albicans* | Transcriptome analysis uncovers a link between copper metabolism, and both fungal fitness and antifungal sensitivity in the opportunistic yeast *Candida albicans* | Khemiri I, Tebbji F, Sellam A | PMID: 32508775 |
| GSE154488 | *C. albicans* | Global Translational Landscape of the *Candida albicans* Morphological Transition | Kadosh D, Choudhary S | PMID: 33585865 |
| GSE173668 | *C. albicans* | The Negative Effect of Protein Phosphatase Z1 Deletion on the Oxidative Stress Tolerance of *Candida albicans* Is Synergistic with Betamethasone Exposure | Pócsi I, Jakab Á | PMID: 34356919 |
| GSE56174 | *C. albicans* | Microevolution of *Candida albicans* in macrophages restores filamentation in a nonfilamentous mutant | Lüttich A, Linde J, Brunke S | PMID: 25474009 |
| GSE68477 | *C. albicans* | Transcriptional landscape of transkingdom communication between *Candida albicans* and *Streptococcus gordonii* | Dutton LC, Paskiewicz KH, Silverman RJ, Splatt PR, Shaw S, Nobbs AH, Lamont RJ, Jenkinson H, Ramsdale M | PMID: 26042999 |
| GSE86540 | *C. albicans* | Effects of pH on the transcriptional profile of adenylate cyclase (cyr1) mutants | Hollomon JM, Grahl N, Koeppen K, Willger SD, Hogan DA | PMID: 27921082 |
| GSE96965 | *C. albicans* | Identification and mode of action of a plant natural product targeting human fungal pathogens | Dorsaz S, Sanglard D | PMID: 28674054 |

| | | | | |
|---|---|---|---|---|
| GSE103952 | *C. difficile* | Deoxycholate induces biofilm formation by *Clostridium difficile* while repressing sporulation and toxin production | Tremblay Y, Monot M | PMID: 31098293 |
| GSE107961 | *C. difficile* | *Clostridium difficile* transcriptome of strain R20291 in response to cysteine by using a time-resolved RNA-sequencing | Wang J, Gu H | PMID: 30172297 |
| GSE63777 | *C. difficile* | Regulation of σK activity and function by SpoIIID during *Clostridium difficile* sporulation | Pishdadian K, Shen A | PMID: 25393584 |
| GSE86152 | *C. difficile* | Effect of Bile Acids on *C. difficile* Growth | Sorg J, Monot M | NA |
| GSE86612 | *C. difficile* | Pathogenomic study of *Clostridium difficile* strains | Monot M, Dupuy B | NA |
| GSE164282 | *F. nucleatum* | Influence of dual-species coaggregation on transcriptional changes of *Streptococcus gordonii* and *Fusobacterium nucleatum* subsp. polymorphum | Liu T, Guo L | PMID: 35071038 |
| GSE106187 | *H. influenza* | RNA SEQ NTHi modA2 | Atack JM, Jennings MP | NA |
| GSE129761 | *H. influenza* | RNA SEQ NTHi modA15 | Atack JM, Jennings MP | PMID: 31320413 |
| GSE129764 | *H. influenza* | RNA SEQ NTHi modA18 | Atack JM, Jennings MP | PMID: 31320413 |
| GSE129786 | *H. influenza* | RNA SEQ NTHi modA10 | Atack JM, Jennings MP | NA |
| GSE131505 | *H. influenza* | Sustained reprioritization of metabolic pathways is associated with persistence and biofilm formation in non-typeable *Haemophilus influenzae* | Harrison A, Hardison RL, Wallace RM, Fitch J, Heimlich DR, O'Bryan M, Sebra RP, Dubois L, St. John-Williams L, White P, Mosely | PMID: 31700653 |

| | | | | |
|---|---|---|---|---|
| | | | M, Thompson J, Justice SS, Mason KM | |
| GSE63900 | *H. influenza* | Dual RNA-sequencing of nontypeable *Haemophilus influenzae* and host cell transcriptomes reveals new aspects of host-pathogen interface | Baddal B, Muzzi A, Censini S, Calogero RA, Torricelli G, Guidotti S, Taddei AR, Covacci A, Pizza M, Rappuoli R, Soriani M, Pezzicoli A | PMID: 26578681 |
| GSE78787 | *M. abscessus* | RNA sequencing of *Mycobacterium abscessus* under infection-relevant conditions | Miranda-CasoLuengo AA, Dinan AM, Loftus BJ | PMID: 27495169 |
| GSE110445 | *P. aeruginosa* | RNA-seq for RpoN | Deng X | NA |
| GSE111903 | *P. aeruginosa* | Sialic acid relieves RetS-dependent inhibition of GacS to induce a chronic infection state of *Pseudomonas aeruginosa* | Wang BX, Cady KC, Ribbeck K, Laub MT | NA |
| GSE112354 | *P. aeruginosa* | Transcriptome analysis of gene expression of PAK-AR2 and 1-9 | You J, Yang H | PMID: 29748556 |
| GSE112597 | *P. aeruginosa* | The *Pseudomonas aeruginosa* PilSR two-component system regulates both twitching and swimming motilities | Kilmury SL, Burrows LL | PMID: 30042200 |
| GSE112911 | *P. aeruginosa* | Genome-wide analysis of Differential Responses of *Pseudomonas aeruginosa* PAO1 and PA14 cells to Interfacial Stress | Niepa TH | PMID: 29259206 |
| GSE121243 | *P. aeruginosa* | RNA-seq for MvfR, RhlR and QscR | Deng X | PMID: 31270321 |
| GSE122048 | *P. aeruginosa* | Transcriptional profiling of *Pseudomonas aeruginosa* and | Tognon M, Köhler T | PMID: 30630428 |

| | | | | |
|---|---|---|---|---|
| | | *Staphylococcus aureus* during in vitro co-culture | | |
| GSE123356 | *P. aeruginosa* | Determination of *Pseudomonas aeruginosa* transcriptome induced by oleic acid or its derived oxylipins 10-HOME and 7,10-DiHOME. | Gadila SK, Martínez E, Campos-Gómez J | PMID: 30793044 |
| GSE124385 | *P. aeruginosa* | Response of *Pseudomonas aeruginosa* to the innate immune system-derived oxidants hypochlorous acid and hypothiocyanous acid | Farrant KV, Spiga L, Davies JC, Williams HD | PMID: 33106346 |
| GSE125646 | *P. aeruginosa* | Transcriptional modulation of Quoring-sensing signaling throught QslA in *Pseudomonas aeruginosa* PA14 and PAO1. | Lomas R, Conesa A, Bleves S, Voulhoux R, Ize B, Gimenez M, Laubier A, Soscia C, Chauvet C | NA |
| GSE130190 | *P. aeruginosa* | Genomic, transcriptomic, and structural analysis of *Pseudomonas* virus PA5oct highlights the molecular complexity among Jumbo phages | Blasdel B, Lood C, Danis-Wlodarczyk K | PMID: 32154616 |
| GSE136111 | *P. aeruginosa* | Conditional Hfq association with small non-coding RNAs in *Pseudomonas aeruginosa* revealed through comparative UV crosslinking immunoprecipitation followed by high-throughput sequencing [RNA-Seq] | Chihara K, Bischler T, Barquist L, Monzon V, Noda N, Vogel J, Tsuneda S | PMID: 31796567 |
| GSE138731 | *P. aeruginosa* | Next Generation Sequencing Facilitates Quantitative Analysis | Li W, Li H | NA |

| | | | | |
|---|---|---|---|---|
| | | of control of *Pseudomonas aeruginosa* PAO1 and farnesol-treated P. aeruginosa PAO1 Transcriptomes | | |
| GSE142448 | *P. aeruginosa* | Transcriptome profiles of *P. aeruginosa* treated with manuka honey and its key components | Bouzo D, Cokcetin N, Li L, Ballerin G, Bottomley A, Lazenby J, Whitchurch C, Paulsen I, Hassan K, Harry L | PMID: 32606022 |
| GSE142464 | *P. aeruginosa* | *Pseudomonas aeruginosa* core metabolism exerts a widespread growth-independent control on virulence | Promponas VJ, Apidianakis Y | PMID: 32528034 |
| GSE148597 | *P. aeruginosa* | Transcriptional profiling of *Pseudomonas aeruginosa* and *Candida albicans* grown in mono-cultures and co-culture. | Doing G, Hogan DA | PMID: 32813693 |
| GSE148955 | *P. aeruginosa* | RNA-Seq analysis of *Pseudomonas aeruginosa* response to testosterone | Vidaillac C | PMID: 32994320 |
| GSE152480 | *P. aeruginosa* | Control of a programmed cell death pathway in *Pseudomonas aeruginosa* by a ppGpp-responsive antiterminator [RNAtag-seq] | Pena JM, McFarland KA, Kambara TK, Ramsey KM, Deighan P, Dove SL | PMID: 33731715 |
| GSE153067 | *P. aeruginosa* | Introducing Differential RNAseq mapping to track the phage infection process for *Pseudomonas* virus ϕKZ | Wicke L, Ponath F, Coppens L, Gerovac M, Lavigne R, Vogel J | PMID: 33103565 |

| | | | | |
|---|---|---|---|---|
| GSE156995 | *P. aeruginosa* | Transcriptomic profiles of *Pseudomonas aeruginosa* PA14 grown in the presence or absence of mucins or mucin-glycans | Ribbeck K, Wheeler K, Wang B | PMID: 33125866 |
| GSE163234 | *P. aeruginosa* | The LysR-type transcriptional regulator BsrA (PA2121) is engaged in the control of vital metabolic pathways in *Pseudomonas aeruginosa* (RNA-seq) | Modrzejewska M, Kawalek A, Bartosik AA | PMID: 34254827 |
| GSE163248 | *P. aeruginosa* | Dkstatin-treated *Pseudomonas* aeruginosa trascriptome | Min KB, Yoon SS | NA |
| GSE166602 | *P. aeruginosa* | Targeting bacterial gyrase with cystobactamid, fluoroquinolone and aminocoumarin antibiotics induces distinct molecular signatures in *Pseudomonas aeruginosa* | Franke R, Overwin H, Häussler S, Brönstrup M | PMID: 34254824 |
| GSE166986 | *P. aeruginosa* | The DEAD-box RNA helicases RhlE2 is a global regulator of *Pseudomonas aeruginosa* lifestyle and pathogenesis | Hausmann S, Gonzalez D, Geiser J, Valentini M | PMID: 34151378 |
| GSE179150 | *P. aeruginosa* | Comparison of the *Pseudomonas aeruginosa* PA2504 deficient strain and the PAO1161 wild type strain | Drabinska J, Kraszewska E | PMID: 34575996 |
| GSE71880 | *P. aeruginosa* | Transcriptome analyses of continuous cultures of *Pseudomonas aeruginosa* growing slowly under oxygen and nitrate limitation | Thao S, Harwood CS, Oda Y, Hager KR, Radey M, Brittnacher MJ, Hayden HS | NA |

| | | | | |
|---|---|---|---|---|
| GSE87213 | *P. aeruginosa* | Differential gene expression of *Pseudomonas aeruginosa* ΔPA14_22470 (ΔPA3225) versus UCBPP-PA14 wildtype in planktonic and biofilm cells | Hall CW, Zhang L, Mah T | PMID: 28584154 |
| GSE99729 | *P. aeruginosa* | Transcriptional and mutational profiling of an aminoglycoside resistant *Pseudomonas aeruginosa* small colony variant | Schniederjans M, Koska M, Häussler S | NA |
| GSE99981 | *P. aeruginosa* | Next generation sequencing of *Pseudomonas aeruginosa* PAO1 in the presence of Copper Nanoparticles Stress | Guo J, Gao S | NA |
| GSE124206 | *Porphyromonas* species | Comparison of *Porphyromonas gingivalis* W83 gene expression when grown with or without galactose | Moye ZD, Gormley CM, Davey ME | PMID: 30552185 |
| GSE78126 | *Porphyromonas* species | *Porphyromonas gingivalis* transcriptome in presence or absence of pABA | Kuboniwa M, Hendrickson EL, Wang Q, Sakanala A, Wang T, Beck DA, Miller DP, Whiteley M, Amano A, Wang H, Hackett M, Lamont RJ | PMID: 28293219 |
| GSE106456 | *S. aureus* | The conserved regulatory RNA RsaE down-regulates the arginine degradation pathway in *Staphylococcus aureus* [ssRNA-Seq] | Rochat T, Gautheret D, Toffano-Nioche C, Bouloc P | PMID: 29986060 |
| GSE122048 | *S. aureus* | Transcriptional profiling of *Pseudomonas* | Tognon M, Köhler T | PMID: 30630428 |

| | | | | |
|---|---|---|---|---|
| | | *aeruginosa* and *Staphylococcus aureus* during in vitro co-culture | | |
| GSE122065 | *S. aureus* | MS2-affinity purification coupled with RNA sequencing (MAPS) reveals *S. aureus* RsaI sRNA targetome | Caldelari I, Marzi S, Romby P | NA |
| GSE125741 | *S. aureus* | Effect of apicidin on gene expression in *Staphylococcus aureus* USA300 strain LAC | Parlet CP, Kavanaugh JS, Crosby HA, Raja HA, El-Elimat T, Todd DA, Pearce CJ, Cech NB, Oberlies NH, Horswill AR | PMID: 30943400 |
| GSE130777 | *S. aureus* | Regulation of gene expression by MgrA in *Staphylococcus aureus* USA300 strain LAC | Crosby HA, Horswill AR | PMID: 27144398 |
| GSE132179 | *S. aureus* | RNA-Seq of WT *Staphylococcus aureus* and dpurR grown to exponential and stationary phase | Sause WE, Torres VJ | PMID: 31217288 |
| GSE138090 | *S. aureus* | Potentiating the activity of berberine for *Staphylococcus aureus* in a combinatorial treatment with thymol | Aksoy CS, Sariyar-Akbulut B | PMID: 33010366 |
| GSE139071 | *S. aureus* | Transcriptome Comparison of *Staphylococcus aureus* phoU Homologies: Genes Deletion strain and the Parent Strain in stationary phase (12 h) | Shang Y, Wang X, Qu D | PMID: 32670206 |
| GSE139659 | *S. aureus* | Intracellular *Staphylococcus aureus* persisters upon antibiotic exposure | Peyrusson F, Varet H, Legendre R, Sismeiro O, Coppée J, Wolz C, Tenson T, Van Bambeke F | PMID: 32366839 |
| GSE147157 | *S. aureus* | RNA-seq analysis of active anthraquinones | Song Z | PMID: 34539607 |

| | | | | |
|---|---|---|---|---|
| | | as anti-biofilm agents of methicillin-resistant *Staphylococcus aureus* | | |
| GSE148024 | *S. aureus* | Molecular reprogramming and phenotype switching in *Staphylococcus aureus* lead to high antibiotic persistence and affect therapy success | Huemer M, Mairpady Shambat S | PMID: 33574060 |
| GSE152833 | *S. aureus* | Bioplatforms Australia: Antibiotic Resistant Sepsis Pathogens Framework Initiative - *Staphylococcus aureus* BPH2986 | Not Listed | NA |
| GSE152834 | *S. aureus* | Bioplatforms Australia: Antibiotic Resistant Sepsis Pathogens Framework Initiative - *Staphylococcus aureus* BPH2947 | Not Listed | NA |
| GSE152835 | *S. aureus* | Bioplatforms Australia: Antibiotic Resistant Sepsis Pathogens Framework Initiative - *Staphylococcus aureus* BPH2900 | Not Listed | NA |
| GSE152837 | *S. aureus* | Bioplatforms Australia: Antibiotic Resistant Sepsis Pathogens Framework Initiative - *Staphylococcus aureus* BPH2819 | Not Listed | NA |
| GSE152838 | *S. aureus* | Bioplatforms Australia: Antibiotic Resistant Sepsis Pathogens Framework Initiative - *Staphylococcus aureus* BPH2760 | Not Listed | NA |
| GSE40864 | *S. aureus* | Investigating the sRNA and mRNA transcriptional response to antibiotics in methicillin-resistant | Howden BP | PMID: 23733475 |

| | | | | |
|---|---|---|---|---|
| | | *Staphylococcus aureus* using Illumina RNAseq | | |
| GSE56294 | *S. aureus* | Comparative genomics of *Staphylococcus aureus* and the application of a “pan-genome” for assigning RNA-Seq transcript reads from divergent strains and in vivo human nasal samples | Chaves-Moreno D, Wos-Oxley ML, Jáuregui R, Medina E, Pieper DH, Oxley AP | PMID: 26717500 |
| GSE68772 | *S. aureus* | The C-terminal region of the RNA helicase CshA is required for the interaction with the degradosome and turnover of bulk RNA in the opportunistic pathogen *Staphylococcus aureus*. | Giraud C, Hausmann S, Lemeille S, Prados J, Redder P, Linder P | PMID: 25997461 |
| GSE83995 | *S. aureus* | Characterization of the LFR Genomic Islet in *Staphylococcus aureus* CC30 | Gill SR | NA |
| GSE89791 | *S. aureus* | Changes in relative transcript amounts caused by ΔftsH in *Staphylococcus aureus* USA300 | Taeok B, Rusch DB, Ford J | PMID: 28814746 |
| *Stenotrophomonas maltophilia* GSE99563 | *S. aureus* | The *Staphylococcus aureus* α-Acetolactate Synthase ALS Confers Resistance to Nitrosative Stress | Carvalho SM, de Jong A, Kloosterman TG, Kuipers OP, Saraiva LM | PMID: 28744267 |
| GSE121347 | *S. maltophilia* | FLR19 response to pH | Gallagher T, Whiteson K | NA |
| GSE125704 | *S. maltophilia* | *Stenotrophomonas maltophilia* differential gene expression in synthetic cystic fibrosis sputum identifies shared and cystic fibrosis strain-specific expression responses to the sputum environment | Willsey GG, Eckstrom K, LaBauve AE, Hinkel LA, Schutz K, LiPuma JJ, Wargo MJ | PMID: 31109991 |

| | | | | |
|---|---|---|---|---|
| GSE141276 | *S. maltophilia* | The inactivation of enzymes belonging to the central carbon metabolism, a novel mechanism of developing antibiotic resistance | Gil-Gil T, Corona F, Martínez JL, Bernardini A | PMID: 32487742 |
| GSE101583 | *Streptococcus* species | KhpA and KhpB Regulate Peptidoglycan Synthesis and Cell Division in the "Superbug" *Streptococcus pneumoniae* | Zheng JJ, Tsui HT, Winkler ME | PMID: 28941257 |
| GSE109680 | *Streptococcus* species | A vaginal tract signal detected by the GBS SaeRS system elicits transcriptomic changes and enhances murine colonization. | Cook LC, Federle MJ, Hu H, Maienschein-Cline M | PMID: 29378799 |
| GSE120640 | *Streptococcus* species | Molecular singularities in a pheromone sensor triumvirate desynchronize the competence-predation interplay in the human commensal *Streptococcus salivarius* | Mignolet J, Cerckel G, Damoczi J, Ledesma-Garcia L, Coenye T, Hols P | PMID: 31433299 |
| GSE132611 | *Streptococcus* species | RNA-Sequencing of *S. pneumoniae* when exposed to the bacteriophage SpSL1 | Furi L, Crawford LA, Oggioni MR | PMID: 31285240 |
| GSE132733 | *Streptococcus* species | Identifying the regulon of Two Component System 7 (SPD_0157 and SPD_0158) of *Streptococcus pneumoniae* | Andreassen PR, Pakula K, Jørgensen MG | PMID: 32130269 |
| GSE139093 | *Streptococcus* species | The impact of depletion of iron or manganese on the transcriptome of *Streptococcus mutans* | Kajfasz JK, Ganguly T, Lemos J | PMID: 31915219 |
| GSE142362 | *Streptococcus* species | Transcriptomic comparison of D39 WT and SPD_0588 deletion | Zhang C | NA |

| | | | | |
|---|---|---|---|---|
| | | mutant in *Streptococcus pneumoniae* D39 with 1 µg/ml methionine | | |
| GSE142458 | *Streptococcus* species | Global gene expression analysis of MGAS10870 serotype of Group A *Streptococcus* (*Streptococcus pyogenes*/GAS) in the absence or presence of 125µg/ml human calprotectin protein | Makthal N, Kumaraswami M | NA |
| GSE148401 | *Streptococcus* species | A dedicated three component regulatory system (GSP/Ghk/Grr) activates gallocin transcription in *Streptococcus gallolyticus* UCN34 | Proutière A, Varet H, Legendre R, Sismeiro O, Dramsi S | PMID: 33402539 |
| GSE149546 | *Streptococcus* species | S1-Domain RNA Binding Protein (CvfD) Is a New Post-Transcriptional Regulator That Mediates Cold Shock, Phosphate Transport, and Virulence in *Streptococcus pneumoniae* D39 | Rusch DB, Winkler ME | NA |
| GSE152821 | *Streptococcus* species | Bioplatforms Australia: Antibiotic Resistant Sepsis Pathogens Framework Initiative - *Streptococcus pyogenes* SP444 | Not Listed | NA |
| GSE152822 | *Streptococcus* species | Bioplatforms Australia: Antibiotic Resistant Sepsis Pathogens Framework Initiative - *Streptococcus pyogenes* PS006 | Not Listed | NA |
| GSE152823 | *Streptococcus* species | Bioplatforms Australia: Antibiotic Resistant Sepsis Pathogens | Not Listed | NA |

| | | | | |
|---|---|---|---|---|
| | | Framework Initiative - *Streptococcus pyogenes* PS003 | | |
| GSE152824 | *Streptococcus* species | Bioplatforms Australia: Antibiotic Resistant Sepsis Pathogens Framework Initiative - *Streptococcus pyogenes* HKU419 | Not Listed | NA |
| GSE152826 | *Streptococcus* species | Bioplatforms Australia: Antibiotic Resistant Sepsis Pathogens Framework Initiative - *Streptococcus pyogenes* 5448 | Not Listed | NA |
| GSE152827 | *Streptococcus* species | Bioplatforms Australia: Antibiotic Resistant Sepsis Pathogens Framework Initiative - *Streptococcus pneumoniae* 4559 | Not Listed | NA |
| GSE152828 | *Streptococcus* species | Bioplatforms Australia: Antibiotic Resistant Sepsis Pathogens Framework Initiative - *Streptococcus pneumoniae* 4496 | Not Listed | NA |
| GSE152829 | *Streptococcus* species | Bioplatforms Australia: Antibiotic Resistant Sepsis Pathogens Framework Initiative - *Streptococcus pneumoniae* 180/2 | Not Listed | NA |
| GSE152832 | *Streptococcus* species | Bioplatforms Australia: Antibiotic Resistant Sepsis Pathogens Framework Initiative - *Streptococcus pneumoniae* 180/15 | Not Listed | NA |
| GSE152965 | *Streptococcus* species | Bioplatforms Australia: Antibiotic Resistant Sepsis Pathogens Framework Initiative - *Streptococcus pneumoniae* 947 | Not Listed | NA |

| | | | | |
|---|---|---|---|---|
| GSE153766 | *Streptococcus* species | Transcriptomic analysis of *Streptococcus suis* in response to ferrous iron and cobalt toxicity | Zheng C, Jia M, Wei M, Zhang Y | PMID: 32887434 |
| GSE158512 | *Streptococcus* species | The CovR environmental sensor orchestrates competence bimodality in salivarius streptococci | Knoops A, Vande Capelle F, Verhaegen M, Mignolet J, Fontaine L, Goffin P, Sass A, Coenye T, Delvigne F, Ledesma-Garcia L, Hols P | PMID: 35089064 |
| GSE160737 | *Streptococcus* species | Transcriptome comparison between emergent and historical emm4 group A *Streptococcus* strains from North America | DebRoy S, Sanson MA, Odo C, Shelburne SA, Flores AR | NA |
| GSE164206 | *Streptococcus* species | Transcriptomic analysis of *Streptococcus suis* in response to excess manganese | Zheng C, Wei M, Qiu J, Jia M | NA |
| GSE164207 | *Streptococcus* species | RNA sequencing analysis of the genes regulated by TroR in *Streptococcus suis* | Zheng C, Wei M, Qiu J, Jia M | NA |
| GSE165796 | *Streptococcus* species | Transcriptome analysis of *Streptococcus mutans* UA159 and SMU_1995c knock out strain | Pan Y, Li Y, Zou J | NA |
| GSE167217 | *Streptococcus* species | Improved transformation efficiency of group A *Streptococcus* by inactivation of a type I restriction modification system | Finn MB, Ramsey KM, Dove SL, Wessels MR | PMID: 33914767 |
| GSE167895 | *Streptococcus* species | RNA sequencing analysis of *Streptococcus agalactiae* 874391 (wild-type) during Cu intoxication | Ulett GC | NA |

| | | | | |
|---|---|---|---|---|
| GSE167896 | *Streptococcus* species | RNA sequencing analysis of CopY-deficient *Streptococcus agalactiae* | Ulett GC | NA |
| GSE167897 | *Streptococcus* species | RNA sequencing analysis of CopY-deficient *Streptococcus agalactiae* during Cu stress | Ulett GC | NA |
| GSE167898 | *Streptococcus* species | RNA sequencing analysis of covR-deficient *Streptococcus agalactiae* | Ulett GC | NA |
| GSE167899 | *Streptococcus* species | RNA sequencing analysis of covR-deficient *Streptococcus agalactiae* during Zn intoxication | Ulett GC | NA |
| GSE167900 | *Streptococcus* species | RNA sequencing analysis of covR-deficient *Streptococcus agalactiae* during Cu intoxication | Ulett GC | NA |
| GSE167901 | *Streptococcus* species | RNA sequencing analysis of SczA-deficient *Streptococcus agalactiae* | Ulett GC | NA |
| GSE167902 | *Streptococcus* species | RNA sequencing analysis of SczA-deficient *Streptococcus agalactiae* during Zn intoxication | Ulett GC | NA |
| GSE167903 | *Streptococcus* species | RNA sequencing analysis of SczA-deficient *Streptococcus agalactiae* during Cu intoxication | Ulett GC | NA |
| GSE173362 | *Streptococcus* species | Identification of group A *Streptococcus* genes directly regulated by CsrRS and novel intermediate regulators [RNA-seq] | Finn MB, Ramsey KM, Dove SL, Wessels MR | PMID: 34253064 |
| GSE181516 | *Streptococcus* species | The LiaFSR transcriptome reveals an interconnected | Sanson MA, Vega LA, Shah B, Regmi S, Cubria MB, | PMID: 34370508 |

| | | | | |
|---|---|---|---|---|
| | | regulatory network in group A *Streptococcus* | Horstmann N, Shelburne III SA, Flores AR | |
| GSE54199 | *Streptococcus* species | Transcriptome and marker frequency analysis of *E. coli* (control vs. Trimethoprim) and *S. pneumoniae* (control vs. HPUra/Kanamycin) | Slager J, Kjos M, Attaiech L, Veening JW | PMID: 24725406 |
| GSE67533 | *Streptococcus* species | Transcriptome study of the covS-regulation of a Group A *Streptococcus pyogenes* M23ND | Liang Z, Bao Y, Mayfield JA, Ploplis VA, Castellino FJ | PMID: 26216843, PMID: 27329479 |
| GSE68477 | *Streptococcus* species | Transcriptional landscape of transkingdom communication between *Candida albicans* and *Streptococcus gordonii* | Dutton LC, Paskiewicz KH, Silverman RJ, Splatt PR, Shaw S, Nobbs AH, Lamont RJ, Jenkinson H, Ramsdale M | PMID: 26042999 |
| GSE77021 | *Streptococcus* species | Changes in relative transcript amounts caused by Δpbp1a, double Δpbp1a mltG(Δ488bp), and triple Δpbp1a mltG(Δ488bp) Δpbp2b mutations in *Streptococcus pneumoniae* | Tsui HT, Zimmer K, Rusch DB, Bruce K, Winkler ME | PMID: 26933838 |
| GSE86854 | *Streptococcus* species | Whole genome transcriptome analysis of serotype M1 Group A *Streptococcus* (*Streptococcus pyogenes*) wild type and isogenic FabT mutants grown at 35C and 40C. | Musser JM, Beres SB, Eraso J | PMID: 27600505 |
| GSE87759 | *Streptococcus* species | RNA-seq analysis of genes regulated by stk in *Streptococcus suis* serotype 2 | Zhang C, Zhou R, Li L | PMID: 28326294 |

| | | | | |
|---|---|---|---|---|
| GSE89964 | *Streptococcus* species | Analysis of differential gene expression in the *Streptococcus sanguinis* brpT mutant | Stone VN, Xu P | PMID: 28046010 |
| GSE90021 | *Streptococcus* species | Analysis of differential gene expression in the *Streptococcus sanguinis* SSA_0351 mutant | El Rami FE, Xu P | PMID: 28869408 |
| GSE95709 | *Streptococcus* species | Transcriptomic comparison of wild-type and ptvR deletion mutant of *Streptococcus pneumoniae* D39 | Liu X, Li J, Feng Z, Luo Y, Veening J, Zhang J | PMID: 28484041 |
| GSE97218 | *Streptococcus* species | Analysis of differential gene expression in the *Streptococcus sanguinis* SK36 treated with sub-inhibitory dose of Ampicillin for 3 time periods. | El-Rami FE, Xu P | PMID: 29393020 |
| GSE97357 | *Streptococcus* species | Analysis of differential gene expression in the *Streptococcus sanguinis* SK36 treated with heat shock for 3 periods of time | El-Rami FE, Xu P | NA |

**Table S2. Example Table of Differentially Expressed Genes.** As described in the *A. fumigatus* user story (user story #1), the application was used to compare gene expression for A. fumigatus cultured with P. aeruginosa vs. A. fumigatus cultured alone at a timepoint of 180 minutes. Genes differentially expressed at 180 minutes with p value less than 0.05 were downloaded from the application in a table (514 genes). The resulting list of genes was further filtered to include just those genes with $|\log_2 FC| > 1.5$ (10 genes, shown here).

| Gene ID | $\log_2$FC | $\log_2$CPM | FValue | PValue |
|---|---|---|---|---|
| CADAFUBG00000399 | -1.90 | 0.44654893 | 10.5065718 | 0.00359435 |
| CADAFUBG00004661 | 2.11 | -0.2982177 | 6.29919199 | 0.0195418 |
| CADAFUBG00007159 | 2.15 | -0.3214534 | 6.56882317 | 0.02079987 |
| CADAFUBG00007744 | 1.60 | 1.07297809 | 4.33649208 | 0.04858159 |
| CADAFUBG00007752 | 1.74 | -0.0079859 | 4.9528957 | 0.0361036 |
| CADAFUBG00007840 | -2.04 | 0.73791847 | 8.90182797 | 0.00662776 |
| CADAFUBG00008551 | 2.05 | -0.2801763 | 5.66923059 | 0.02847043 |
| CADAFUBG00008638 | 2.47 | 0.14680563 | 4.38436578 | 0.04745516 |
| CADAFUBG00008721 | -1.53 | 0.29131268 | 6.20741892 | 0.02034956 |
| CADAFUBG00009383 | 1.93 | -0.2967983 | 9.6341365 | 0.00536067 |

**Table S3. Beta Testing Criteria – Filtering Studies.** This table, and the other beta testing tables, we're filled in to identify any bugs in a systematic manner. All identified bugs were subsequently addressed. The app reviewers (three of the paper co-authors, noted in the Contributions section) were also instructed to note suggestions to improve app usability, which were addressed. See the key below the table for species abbreviations (e.g., CA = *Candida albicans*)

| Objective # | Description | Working (Y/N) | Notes | Fixed |
|---|---|---|---|---|
| 1A.1 | For AF studies, do all potential filtering inputs show up before any are selected?<br><br>*Also, make a note here if any of the filtering inputs are repeated, misspelled, or don't make sense to you* | | | |
| 1A.2 | Select each individual filter input (e.g., Strain = A1163) and view the contents of the other drop-down menus. Make sure that they narrow down to reflect the studies matching the selected filter input. After selecting each item, view the filtered table to make sure the proper study pops up. | | | |
| 1A.3 | Select all possible combinations (E.g., Strain = A1163, Treatment = *Pseudomonas* Exposure, Medium = AMM) of filter inputs and ensure that the proper studies materialize when you press the "Finalize Inputs" button | | | |
| 1B.1 | Species: *Bacteroides*<br>See 1A.1 Description | | | |
| 1B.2 | Species: *Bacteroides*<br>See 1A.2 Description | | | |
| 1B.3 | Species: *Bacteroides*<br>See 1A.3 Description | | | |
| 1C.1 | Species: *Burkholderia*<br>See 1A.1 Description | | | |
| 1C.2 | Species: *Burkholderia*<br>See 1A.2 Description | | | |
| 1C.3 | Species: *Burkholderia*<br>See 1A.3 Description | | | |

| | |
|---|---|
| 1D.1 | Species: CA<br>See 1A.1 Description |
| 1D.2 | Species: CA<br>See 1A.2 Description |
| 1D.3 | Species: CA<br>See 1A.3 Description |
| 1E.1 | Species: CD<br>See 1A.1 Description |
| 1E.2 | Species: CD<br>See 1A.2 Description |
| 1E.3 | Species: CD<br>See 1A.3 Description |
| 1F.1 | Species: FN<br>See 1A.1 Description |
| 1F.2 | Species: FN<br>See 1A.2 Description |
| 1F.3 | Species: FN<br>See 1A.3 Description |
| 1G.1 | Species: HI<br>See 1A.1 Description |
| 1G.2 | Species: HI<br>See 1A.2 Description |
| 1G.3 | Species: HI<br>See 1A.3 Description |
| 1H.1 | Species: MA<br>See 1A.1 Description |
| 1H.2 | Species: MA<br>See 1A.2 Description |
| 1H.3 | Species: MA<br>See 1A.3 Description |
| 1I.1 | Species: PA<br>See 1A.1 Description |
| 1I.2 | Species: PA |

| | |
|---|---|
| | See 1A.2 Description |
| 1I.3 | Species: PA<br>See 1A.2 Description |
| 1J.1 | Species: *Porphyromonas*<br>See 1A.1 Description |
| 1J.2 | Species: *Porphyromonas*<br>See 1A.2 Description |
| 1J.3 | Species: *Porphyromonas*<br>See 1A.3 Description |
| 1K.1 | Species: SA<br>See 1A.1 Description |
| 1K.2 | Species: SA<br>See 1A.2 Description |
| 1K.3 | Species: SA<br>See 1A.3 Description |
| 1L.1 | Species: SM<br>See 1A.1 Description |
| 1L.2 | Species: SM<br>See 1A.2 Description |
| 1L.3 | Species: SM<br>See 1A.3 Description |
| 1M.1 | Species: Strep<br>See 1A.1 Description |
| 1M.2 | Species: Strep<br>See 1A.2 Description |
| 1M.3 | Species: Strep<br>See 1A.3 Description |

Key: AF = *Aspergillus fumigatus*, Bacteroides = *Bacteroides* species, Burkholderia = *Burkholderia* species, CA = *Candida albicans*, CD = *Clostridium difficile*, FN = *Fusobacterium nucleatum*, HI = *Haemophilus influenza*, MA = *Mycobacterium abscessus*, PA = *Pseudomonas aeruginosa*, Porphyromonas = *Porphyromonas* species, SA = *Staphylococcus aureus*, SM = *Stenotrophomonas maltophilia*, Strep = *Streptococcus* species

**Table S4. Beta Testing Criteria – Viewing Study Metadata.** This table, and the other beta testing tables, we're filled in to identify any bugs in a systematic manner. All identified bugs were subsequently addressed. The app reviewers (three of the paper co-authors, noted in the Contributions section) were also instructed to note suggestions to improve app usability, which were addressed. See the key below the table for species abbreviations (e.g., CA = *Candida albicans*)

| Objective # | Description | Working (Y/N) | Notes | Fixed |
|---|---|---|---|---|
| 2A.1 | For each AF study, click the 'Get More Info' button and make sure that the correct set of metadata appears | | | |
| 2A.2 | Choose several different sets of filtering inputs – after finalizing each, click the 'Get More Info' button and again make sure that the correct set of metadata appears for each study present. In other words, filtering should not cause the metadata under 'more info' to not match the studies that are shown. | | | |
| 1B.1 | Species: *Bacteroides*<br>See 1A.1 Description | | | |
| 1B.2 | Species: *Bacteroides*<br>See 1A.2 Description | | | |
| 2C.1 | Species: *Burkholderia*<br>See 2A Description | | | |
| 2C.2 | Species: *Burkholderia*<br>See 2A Description | | | |
| 2D.1 | Species: CA<br>See 2A Description | | | |
| 2D.2 | Species: CA<br>See 2A Description | | | |
| 2E.1 | Species: CD<br>See 2A Description | | | |
| 2E.2 | Species: CD<br>See 2A Description | | | |
| 2F.1 | Species: FN<br>See 2A Description | | | |
| 2F.2 | Species: FN<br>See 2A Description | | | |

| | |
|---|---|
| 2G.1 | Species: HI<br>See 2A Description |
| 2G.2 | Species: HI<br>See 2A Description |
| 2H.1 | Species: MA<br>See 2A Description |
| 2H.2 | Species: MA<br>See 2A Description |
| 2I.1 | Species: PA<br>See 2A Description |
| 2I.2 | Species: PA<br>See 2A Description |
| 2J.1 | Species: *Porphyromonas*<br>See 2A Description |
| 2J.2 | Species: *Porphyromonas*<br>See 2A Description |
| 2K.1 | Species: SA<br>See 2A Description |
| 2K.2 | Species: SA<br>See 2A Description |
| 2L.1 | Species: SM<br>See 2A Description |
| 2L.2 | Species: SM<br>See 2A Description |
| 2M.1 | Species: Strep<br>See 2A Description |
| 2M.2 | Species: Strep<br>See 2A Description |

Key: AF = *Aspergillus fumigatus*, Bacteroides = *Bacteroides* species, Burkholderia = *Burkholderia* species, CA = *Candida albicans*, CD = *Clostridium difficile*, FN = *Fusobacterium nucleatum*, HI = *Haemophilus influenza*, MA = *Mycobacterium abscessus*, PA = *Pseudomonas aeruginosa*, Porphyromonas = *Porphyromonas* species, SA = *Staphylococcus aureus*, SM = *Stenotrophomonas maltophilia*, Strep = *Streptococcus* species

**Table S5. Beta Testing Criteria – Running Analysis.** This table, and the other beta testing tables, we're filled in to identify any bugs in a systematic manner. All identified bugs were subsequently addressed. The app reviewers (three of the paper co-authors, noted in the Contributions section) were also instructed to note suggestions to improve app usability, which were addressed. See the key below the table for species abbreviations (e.g., CA = *Candida albicans*)

| Objective # | Description | Working (Y/N) | Notes | Fixed |
|---|---|---|---|---|
| 3A.1 | For each AF study, select all potential comparisons and ensure that a unique differential expression analysis table materializes | | | |
| 3A.2 | For each AF study, select all potential comparisons and ensure that a unique volcano plot materializes | | | |
| 3A.3 | For each AF study, select all potential comparisons and ensure that a unique MA plot materializes | | | |
| 3A.4 | While each comparison is selected, make sure that the 'find a gene' and 'find a pathway' features work properly (i.e., the selected gene is highlighted in red and the selected pathway genes are highlighted in green) for both the volcano plot and the MA plot. Test out a few different genes and pathways to make sure they work | | | |
| 3A.5 | While each comparison is selected, make sure that the data table, volcano plot, and MA plot can all be downloaded, and once they are downloaded, that they appear just as they do in the app window. Test out a few comparisons per study | | | |
| 1B.1 | Species: *Bacteroides*<br>See 1A.1 Description | | | |
| 1B.2 | Species: *Bacteroides*<br>See 1A.2 Description | | | |
| 1B.3 | Species: *Bacteroides*<br>See 1A.3 Description | | | |
| 1B.4 | Species: *Bacteroides*<br>See 1A.2 Description | | | |
| 1B.5 | Species: *Bacteroides*<br>See 1A.3 Description | | | |

| | |
|---|---|
| 3C.1 | Species: *Burkholderia*<br>See 3A Description |
| 3C.2 | Species: *Burkholderia*<br>See 3A Description |
| 3C.3 | Species: *Burkholderia*<br>See 3A Description |
| 3C.4 | Species: *Burkholderia*<br>See 3A Description |
| 3C.5 | Species: *Burkholderia*<br>See 3A Description |
| 3D.1 | Species: CA<br>See 3A Description |
| 3D.2 | Species: CA<br>See 3A Description |
| 3D.3 | Species: CA<br>See 3A Description |
| 3D.4 | Species: CA<br>See 3A Description |
| 3D.5 | Species: CA<br>See 3A Description |
| 3E.1 | Species: CD<br>See 3A Description |
| 3E.2 | Species: CD<br>See 3A Description |
| 3E.3 | Species: CD<br>See 3A Description |
| 3E.4 | Species: CD<br>See 3A Description |
| 3E.5 | Species: CD<br>See 3A Description |
| 3F.1 | Species: FN<br>See 3A Description |
| 3F.2 | Species: FN |

| | |
|---|---|
| | See 3A Description |
| 3F.3 | Species: FN<br>See 3A Description |
| 3F.4 | Species: FN<br>See 3A Description |
| 3F.5 | Species: FN<br>See 3A Description |
| 3G.1 | Species: HI<br>See 3A Description |
| 3G.2 | Species: HI<br>See 3A Description |
| 3G.3 | Species: HI<br>See 3A Description |
| 3G.4 | Species: HI<br>See 3A Description |
| 3G.5 | Species: HI<br>See 3A Description |
| 3H.1 | Species: MA<br>See 3A Description |
| 3H.2 | Species: MA<br>See 3A Description |
| 3H.3 | Species: MA<br>See 3A Description |
| 3H.4 | Species: MA<br>See 3A Description |
| 3H.5 | Species: MA<br>See 3A Description |
| 3I.1 | Species: PA<br>See 3A Description |
| 3I.2 | Species: PA<br>See 3A Description |
| 3I.3 | Species: PA<br>See 3A Description |

| | |
|---|---|
| 3I.4 | Species: PA<br>See 3A Description |
| 3I.5 | Species: PA<br>See 3A Description |
| 3J.1 | Species: *Porphyromonas*<br>See 3A Description |
| 3J.2 | Species: *Porphyromonas*<br>See 3A Description |
| 3J.3 | Species: *Porphyromonas*<br>See 3A Description |
| 3J.4 | Species: *Porphyromonas*<br>See 3A Description |
| 3J.5 | Species: *Porphyromonas*<br>See 3A Description |
| 3K.1 | Species: *Porphyromonas*<br>See 3A Description |
| 3K.2 | Species: SA<br>See 3A Description |
| 3K.3 | Species: SA<br>See 3A Description |
| 3K.4 | Species: SA<br>See 3A Description |
| 3K.5 | Species: SA<br>See 3A Description |
| 3L.1 | Species: SM<br>See 3A Description |
| 3L.2 | Species: SM<br>See 3A Description |
| 3L.3 | Species: SM<br>See 3A Description |
| 3L.4 | Species: SM<br>See 3A Description |
| 3L.5 | Species: SM |

| | |
|---|---|
| | See 3A Description |
| 3M.1 | Species: Strep<br>See 3A Description |
| 3M.2 | Species: Strep<br>See 3A Description |
| 3M.3 | Species: Strep<br>See 3A Description |
| 3M.4 | Species: Strep<br>See 3A Description |
| 3M.5 | Species: Strep<br>See 3A Description |

Key: AF = *Aspergillus fumigatus*, Bacteroides = *Bacteroides* species, Burkholderia = *Burkholderia* species, CA = *Candida albicans*, CD = *Clostridium difficile*, FN = *Fusobacterium nucleatum*, HI = *Haemophilus influenza*, MA = *Mycobacterium abscessus*, PA = *Pseudomonas aeruginosa*, Porphyromonas = *Porphyromonas* species, SA = *Staphylococcus aureus*, SM = *Stenotrophomonas maltophilia*, Strep = *Streptococcus* species

**Figure S1. Zoomed-in image of figure 2 (application workflow), panel 1 (user manual)**

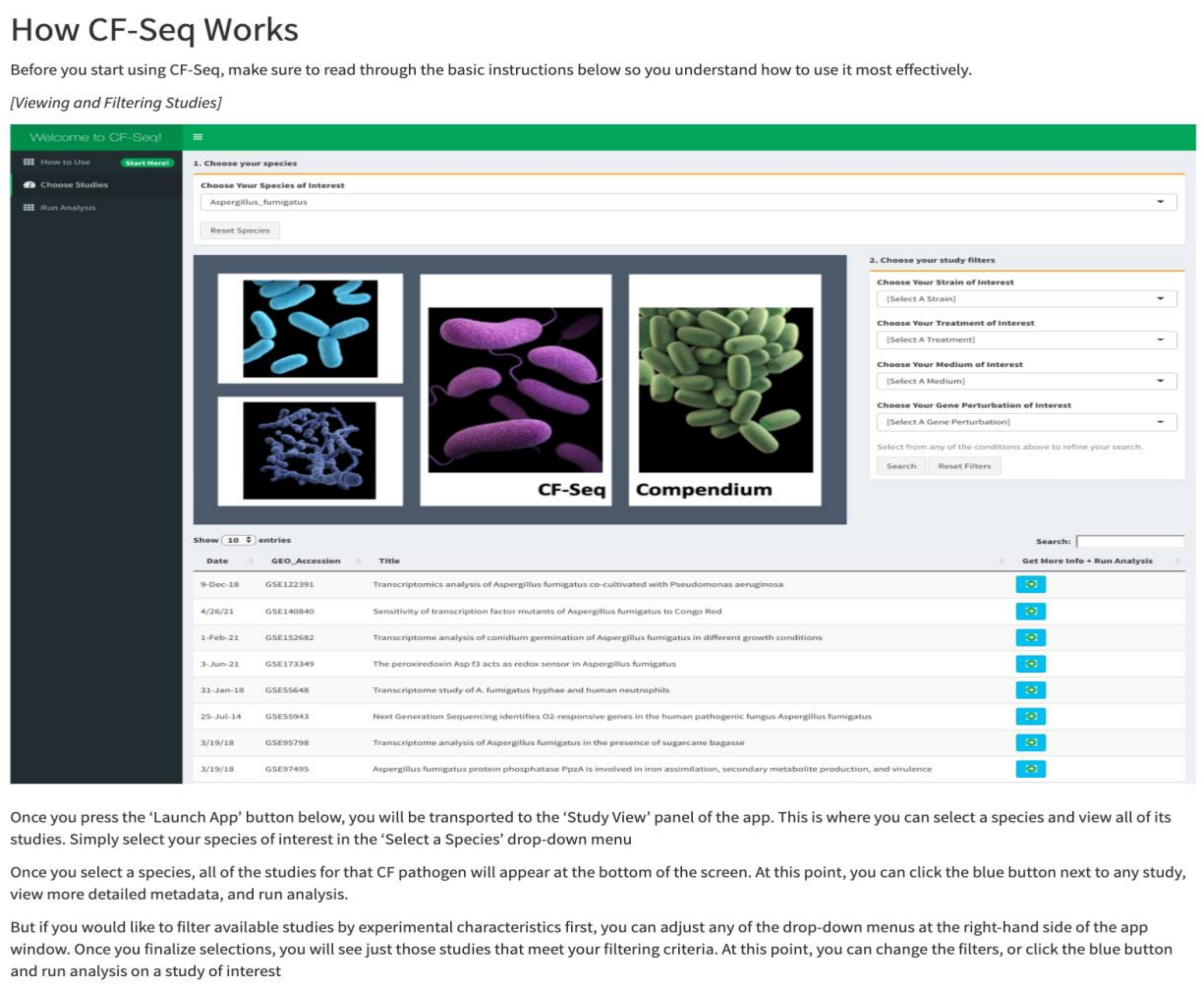

### How CF-Seq Works

Before you start using CF-Seq, make sure to read through the basic instructions below so you understand how to use it most effectively.

*[Viewing and Filtering Studies]*

| Date | GEO_Accession | Title | Get More Info + Run Analysis |
|---|---|---|---|
| 9-Dec-18 | GSE122391 | Transcriptomics analysis of Aspergillus fumigatus co-cultivated with Pseudomonas aeruginosa | |
| 4/26/21 | GSE140840 | Sensitivity of transcription factor mutants of Aspergillus fumigatus to Congo Red | |
| 1-Feb-21 | GSE152682 | Transcriptome analysis of conidium germination of Aspergillus fumigatus in different growth conditions | |
| 3-Jun-21 | GSE173349 | The peroxiredoxin Asp f3 acts as redox sensor in Aspergillus fumigatus | |
| 31-Jan-18 | GSE55648 | Transcriptome study of A. fumigatus hyphae and human neutrophils | |
| 25-Jul-14 | GSE55943 | Next Generation Sequencing identifies O2-responsive genes in the human pathogenic fungus Aspergillus fumigatus | |
| 3/19/18 | GSE95798 | Transcriptome analysis of Aspergillus fumigatus in the presence of sugarcane bagasse | |
| 3/19/18 | GSE97495 | Aspergillus fumigatus protein phosphatase PpzA is involved in iron assimilation, secondary metabolite production, and virulence | |

Once you press the 'Launch App' button below, you will be transported to the 'Study View' panel of the app. This is where you can select a species and view all of its studies. Simply select your species of interest in the 'Select a Species' drop-down menu

Once you select a species, all of the studies for that CF pathogen will appear at the bottom of the screen. At this point, you can click the blue button next to any study, view more detailed metadata, and run analysis.

But if you would like to filter available studies by experimental characteristics first, you can adjust any of the drop-down menus at the right-hand side of the app window. Once you finalize selections, you will see just those studies that meet your filtering criteria. At this point, you can change the filters, or click the blue button and run analysis on a study of interest

**Figure S2. Zoomed-in image of figure 2 (application workflow), panel 2 (study view)**

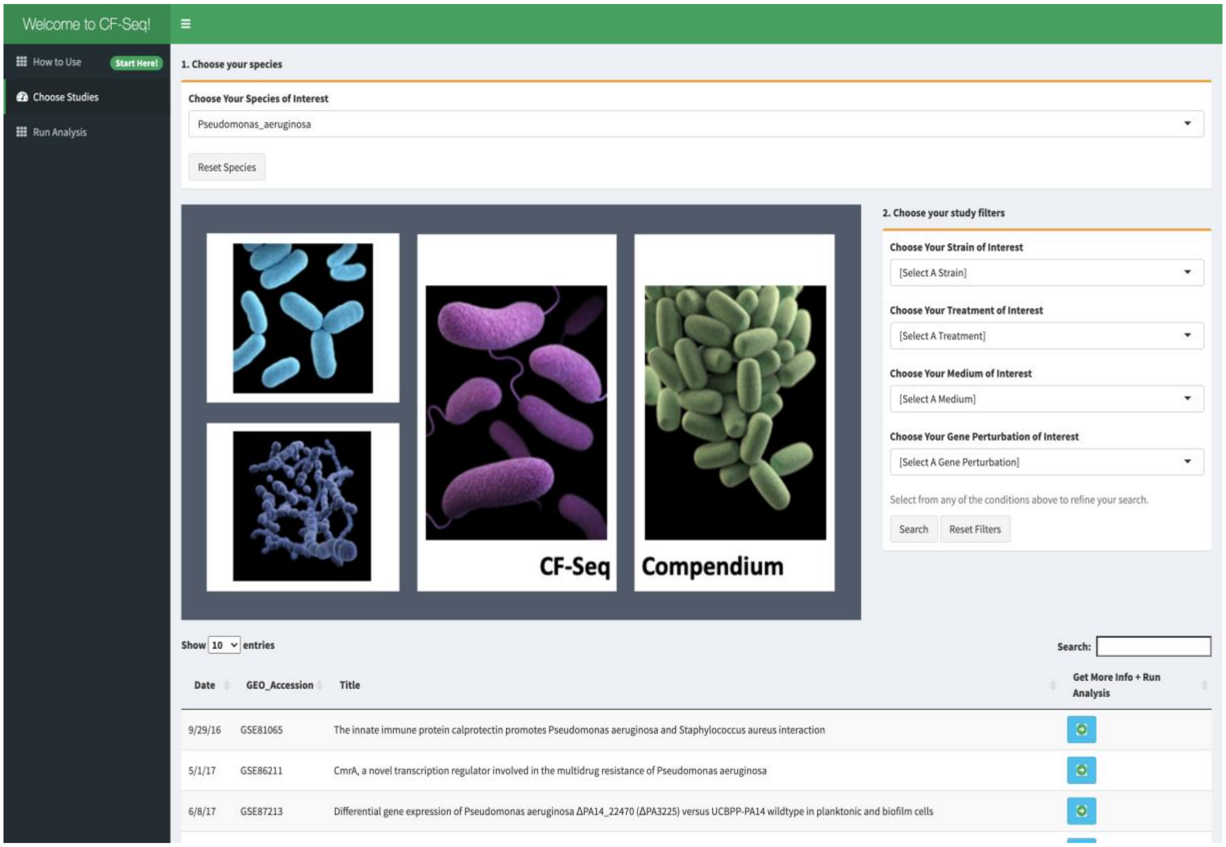

**Figure S3. Zoomed-in image of figure 2 (application workflow), panel 3 (filter studies)**

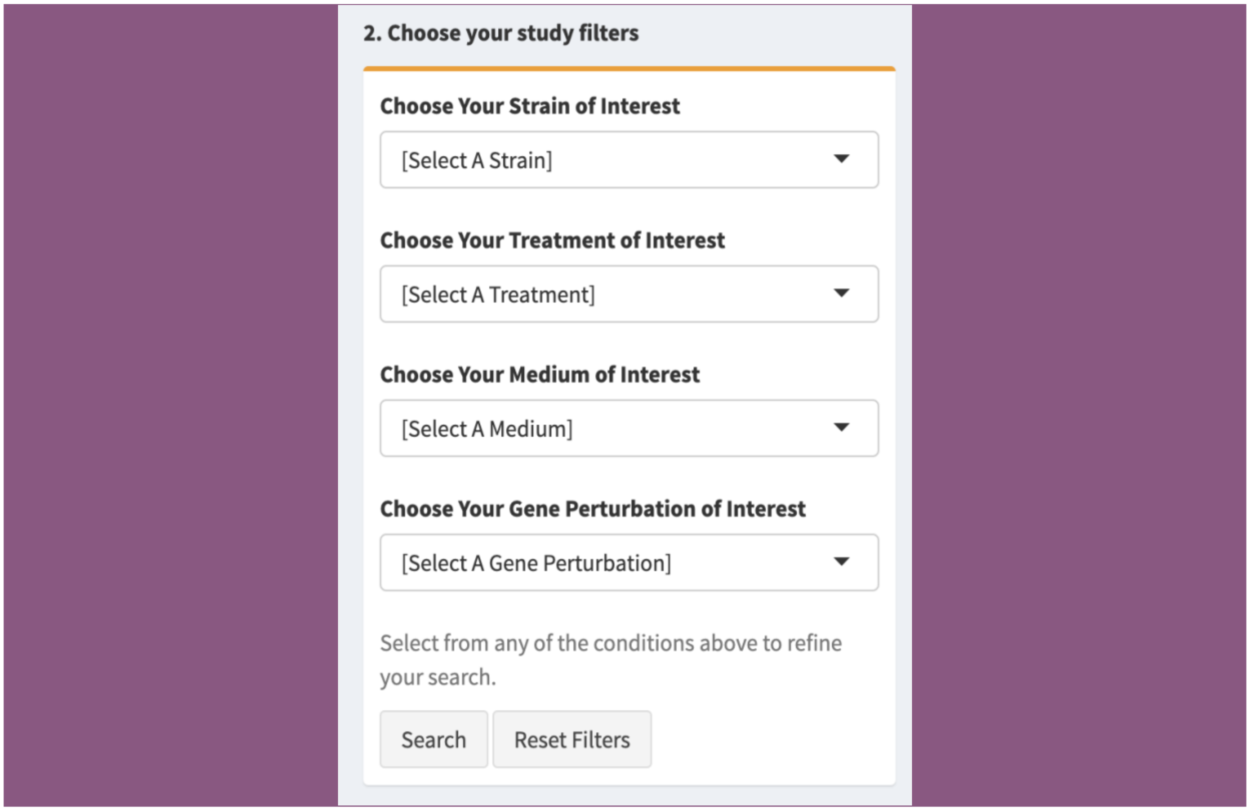

**Figure S4. Zoomed-in image of figure 2 (application workflow), panel 4 (additional metadata)**

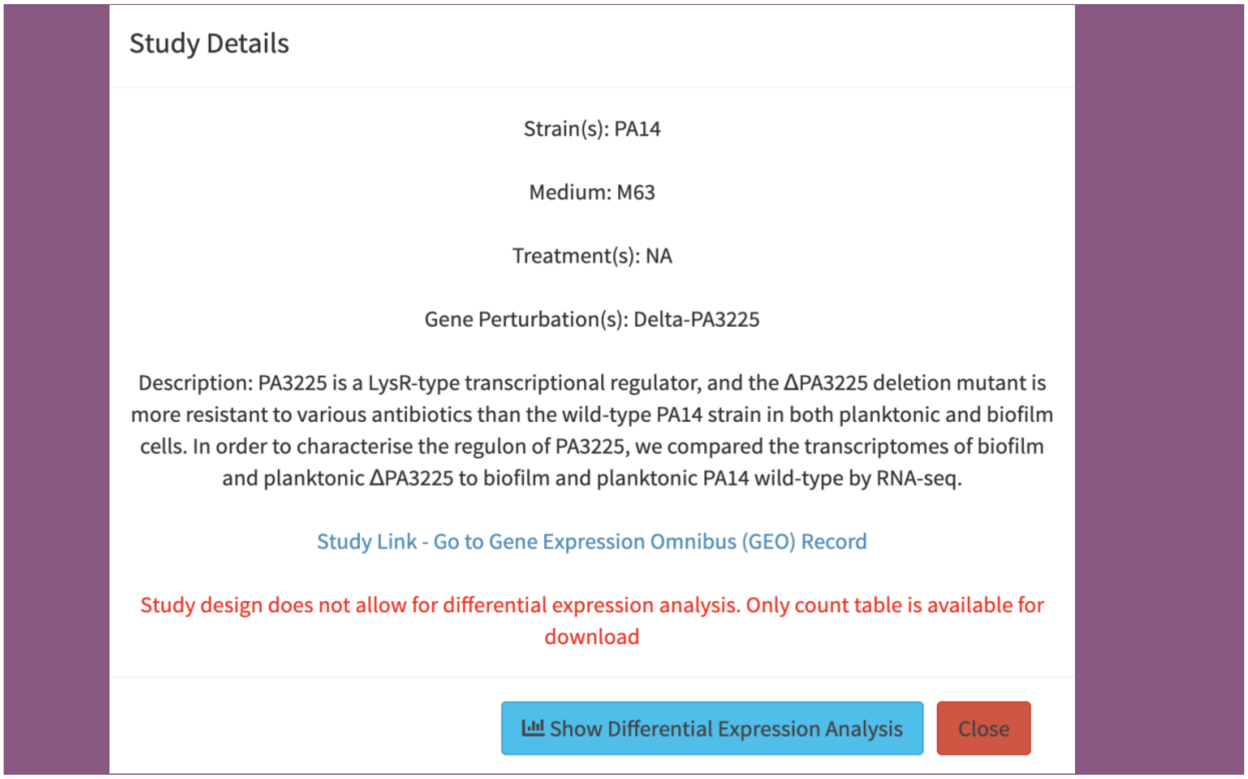

**Figure S5. Zoomed-in image of figure 2 (application workflow), panel 5 (analysis view)**

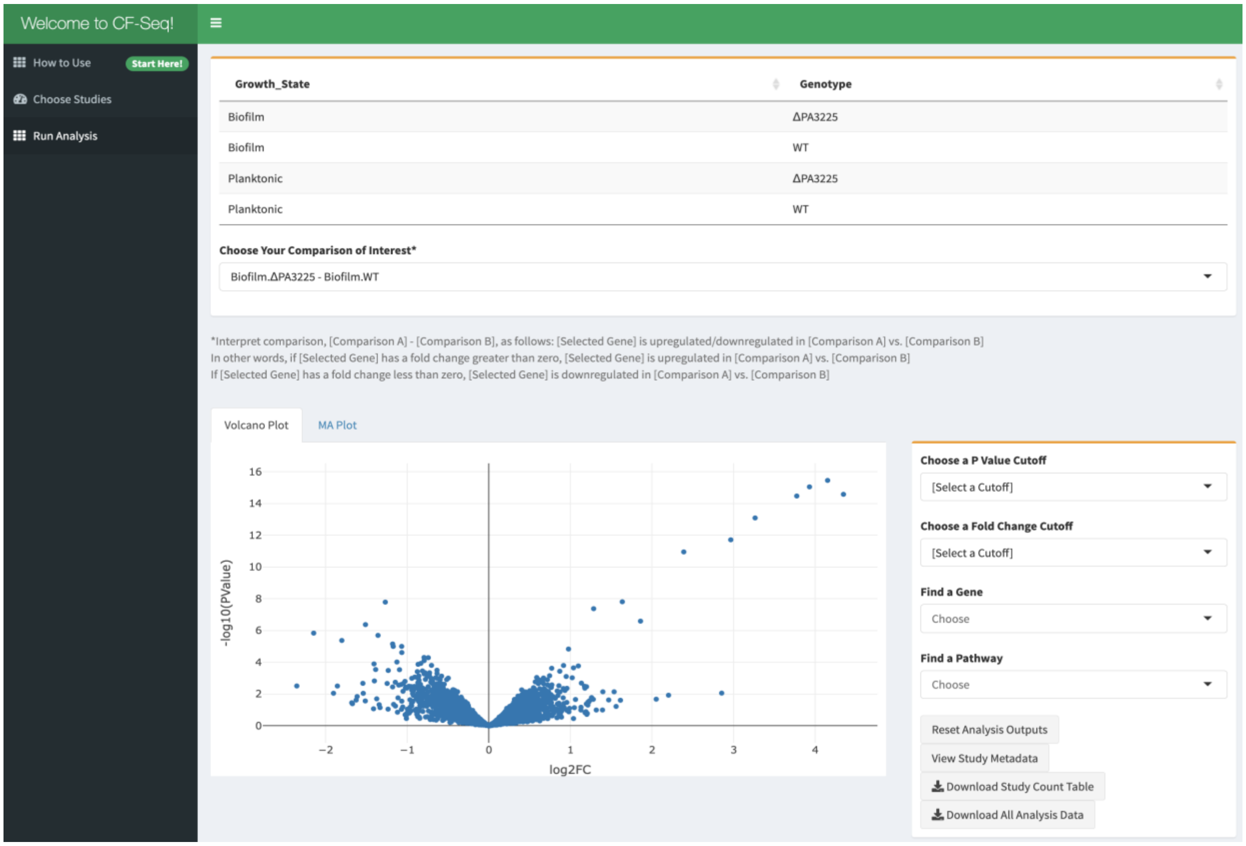

**Figure S6. Zoomed-in image of figure 2 (application workflow), panel 6 (visualize pathway)**

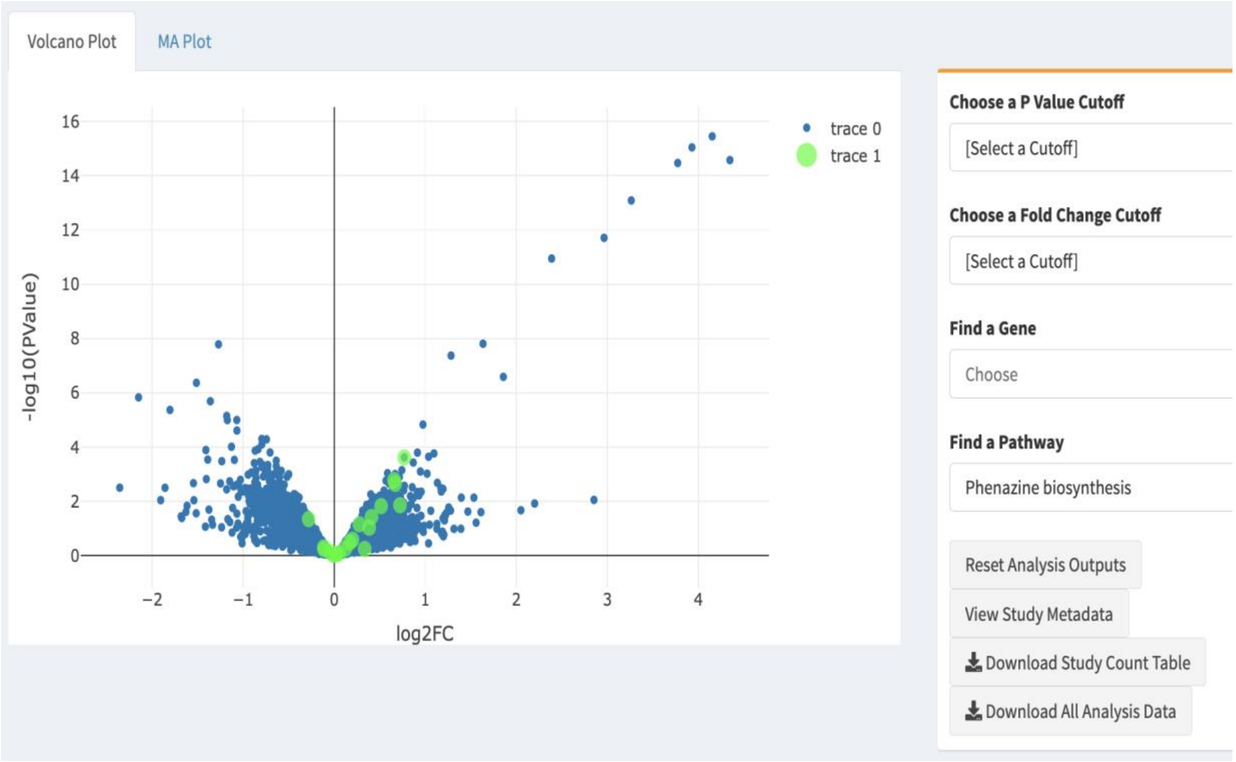
